# Mechanical forces across compartments coordinate cell shape and fate transitions to generate tissue architecture

**DOI:** 10.1101/2022.12.12.519937

**Authors:** Clémentine Villeneuve, Ali Hashmi, Irene Ylivinkka, Elizabeth Lawson-Keister, Yekaterina A. Miroshnikova, Carlos Pérez-González, Bhagwan Yadav, Tao Zhang, Danijela Matic Vignjevic, Marja L. Mikkola, M. Lisa Manning, Sara A. Wickström

**Affiliations:** Stem Cells and Metabolism Research Program, Faculty of Medicine, University of Helsinki, 00290 Helsinki Finland; Department of Cell and Tissue Dynamics, Max Planck Institute for Molecular Biomedicine, 48149 Münster, Germany; Department of Physics and BioInspired Institute, Syracuse University, Syracuse, New York 13244, USA; Cell Biology and Cancer Unit, Institut Curie, PSL Research University, CNRS, Paris, France; School of Chemistry and Chemical Engineering, Shanghai Jiao Tong University, Shanghai 200240, China; Cell and tissue dynamics research program, Institute of Biotechnology, Helsinki Institute of Life Science (HiLIFE), University of Helsinki, Finland; Helsinki Institute of Life Science, Biomedicum Helsinki, University of Helsinki, 00290 Helsinki, Finland; Wihuri Research Institute, Biomedicum Helsinki, University of Helsinki, 00290 Helsinki, Finland

**Keywords:** epithelium, placode, morphogenesis, mechanical force, vertex model

## Abstract

Morphogenesis and cell state transitions must be coordinated in time and space to produce a functional tissue. An excellent paradigm to understand the coupling of these processes is mammalian hair follicle development, initiated by the formation of an epithelial invagination - termed placode – that coincides with the emergence of a designated hair follicle stem cell population. The mechanisms directing the deformation of the epithelium, cell state transitions, and physical compartmentalization of the placode are unknown. Here, we identify a key role for coordinated mechanical forces stemming from contractile, proliferative, and proteolytic activities across the epithelial and mesenchymal compartments in generating the placode structure. A ring of fibroblast cells gradually wraps around the placode cells to generate centripetal contractile forces, which in collaboration with polarized epithelial myosin activity promote elongation and local tissue thickening. These mechanical stresses further enhance and compartmentalize Sox9 expression to promote stem cell positioning. Subsequently, proteolytic remodeling locally softens the basement membrane to facilitate release of pressure on the placode, enabling localized cell divisions, tissue fluidification, and epithelial invagination into the underlying mesenchyme. Together, our experiments and modeling identify dynamic cell shape transformations and tissue-scale mechanical co-operation as key factors for orchestrating organ formation.

## Introduction

The structure of tissues is tightly linked to their function. During formation of functional organs, large-scale changes in tissue elongation, stretching, compression, folding/buckling, and budding impact the shape, position, packing, and contractility state of cells. Conversely, changes in single cell contractility, shape and position locally alter tissue organization and mechanics. Importantly, these tissue and cellscale transformations need to be tightly coordinated in space and time with acquisition of specialized cell states through mechanisms that are poorly understood^1,2^. The formation of the mammalian placode, a lens-shaped multilayered epithelial thickening that gives rise to the hair follicle, serves as an ideal model to study how cell state transitions and morphogenesis are coordinated to generate a specialized tissue structure.

In the mouse embryo, a first wave of periodically spaced pattern of the hair follicle placode arises at late embryonic day 13 (E13) from a homogenous epithelial sheet adhering to a basement membrane that overlays the dermis, a connective tissue containing dispersed mesenchymal cells. This is followed by a second and third wave of morphogenesis around E15.5 and E18, respectively, producing hair follicle placodes morphologically identical and regularly interspersed with the first wave follicle^3,4^. One of the key pathways regulating placode specification is Wnt signaling that acts in a bi-directional, paracrine manner between the epidermis and the underlying dermis. This process commences at E12.5, when secreted epidermal Wnt activates broad mesenchymal β-catenin signaling to initiate the specification of the fibroblast population that later becomes the dermal condensate, an essential structure for hair follicle formation^5^. These fibroblasts, through an unknown signal, then induce hair follicle progenitor fate in the epidermis in patterned preplacodes molecularly identified by Dkk4, Edar, and Fgf20 expression^6–9^. Subsequently, progenitor cell rearrangement and compaction demarcate the physically identifiable placode (Fig. 1a). At the same time, mesenchymal cells migrate and cluster to form a dermal condensate directly underneath the placode^10–12^. Once this spatial pattern has been defined, progenitor cells specify to become follicle cells and activate the expression of additional genes to drive their development into the mature hair follicle that harbors a set of adult stem cells responsible for constant self-renewal of the hair follicle throughout life. One of these genes is Sox9, the master transcription factor required for adult hair follicle stem cells that initially exhibits broader expression but becomes restricted to the placode between E14 and E15, defining the transcriptional profile of future hair follicle stem cells^13,14^. Subsequently the placodes - organized into concentric rings of stem and progenitor cell populations - grow out as longitudinally aligned cylindrical compartments to form the hair follicle 13. However, the physical mechanisms driving the establishment of a Sox9-stem cell population coordinated with stereotypical morphological changes leading to placode invagination are still poorly understood.

**Fig. 1.**
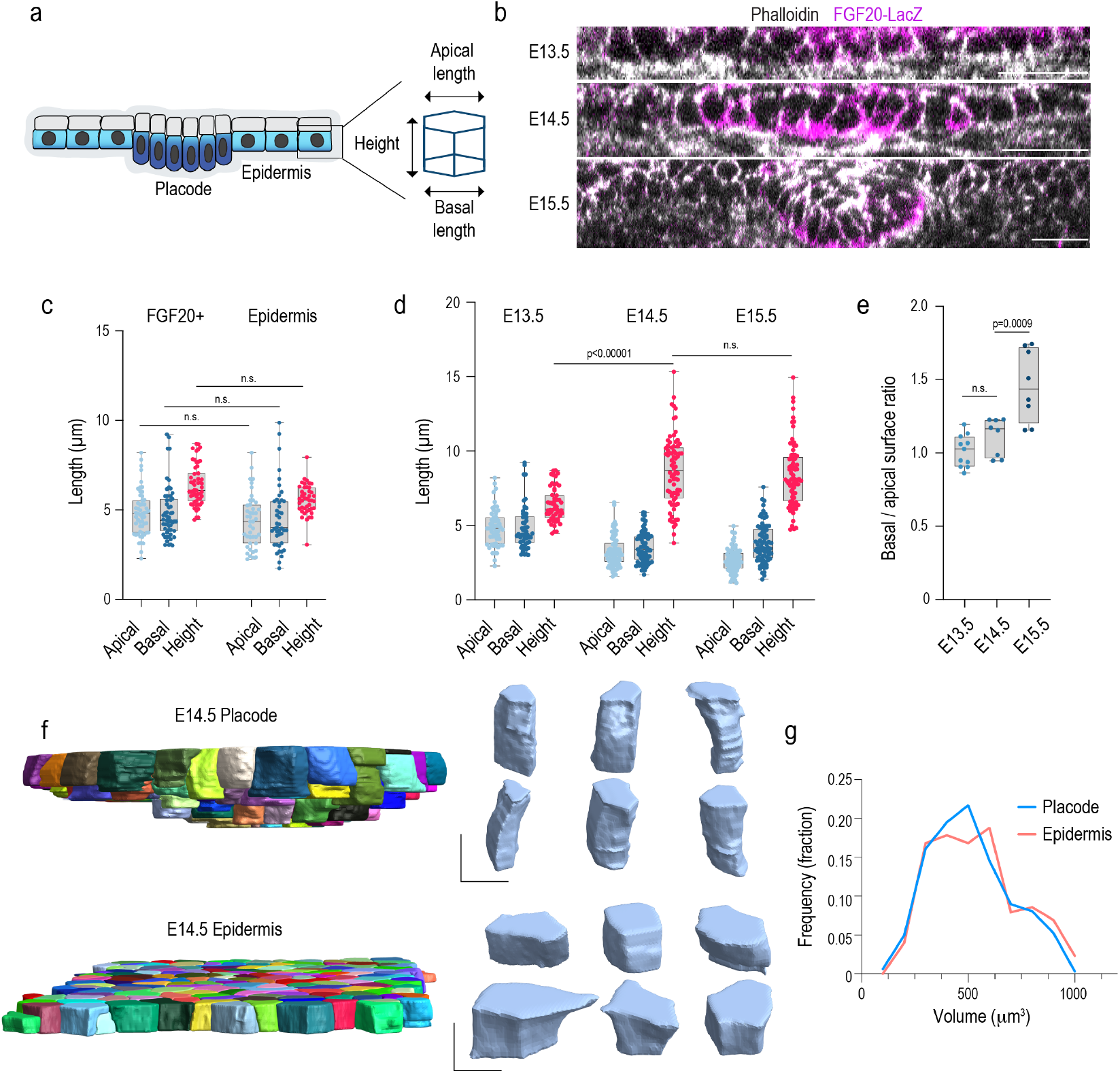
Placode cells undergo isotropic elongation and subsequent basal surface expansion (a) Schematic representation of the epidermis and the morphological parameters quantified. (b) Phalloidin-stained sagittal cross sections of mouse epidermis at embryonic days (E) 13.5-15-5. Scale bars 20 μm. Images representative of 3 mice/group. (c) Quantification of cell axis lengths from placode (marked by FGF20 expression) and epidermis from images in (b). No substantial differences in cell morphology between placode and epidermis are observed (n= 57 cells pooled across 11 placodes/surrounding epidermis from 3 mice). (d) Quantification of cell axis lengths from basal cells of E13.5-E15.5 placodes. Note increased cell height and reduced apical and basal surface lengths in E14.5 and E15.5 placodes and an increase in basal length at E15.5. (n=57(E13.5), 74 (E14.5) and 77 (E15.5) cells from 11 (E13.5), 9 (E14.5) or 8 (E15.5) placodes pooled across 3 mice; Kruskal-Wallis/Dunn’s). (e) Quantification of basal to apical cell surface ratio in E13.5-E15.5 placodes. Note increased basal surface ratio in E15.5 placode (n=11(E13.5), 9 (E14.5) or 8 (E15.5) placodes pooled across 3 mice; Kruskal-Wallis/Dunn’s). (f, g) Quantification and representative images of 3D rendering of cell volumes and shapes from E14.5 placode and epidermis. Note volume-preserving elongation of placode cells. Scale bars 5 μm. (n>250 cells/compartment pooled across 3 mice; p=0.7 Mann-Whitney).

Here we use the murine hair follicle placode as a paradigm to construct and genetically validate a biomechanical model for epithelial sheet transformation into a placode bud. Using whole embryo live imaging, mechanical measurements, genetic manipulations and 3D vertex modeling we show that a combination of epithelial cell-intrinsic actomyosin contractility and extrinsic mechanical stresses from the underlying mesenchyme are required for placode thickening. Importantly, mechanical stress also enhances and spatially restricts Sox9 expression, coupling morphogenetic dynamics to cell type specification and tissue compartmentalization. Subsequent remodeling of the basement membrane facilitates release of accumulated pressure within placodes to induce localized cell divisions necessary for placode budding. Collectively this study identifies a critical role for coordinated tissue- and cell-scale forces across cell compartments in mammalian organogenesis.

## Results

### Placode cells undergo isotropic elongation and subsequent basal surface expansion

Hair follicle placode formation is initiated by epidermal cell rearrangement and compaction to form a physically identifiable thickened structure^10,15^. To investigate morphological changes that generate this specific epithelial deformation, we performed whole mount imaging of mouse epidermis and quantified cell shapes of the basal layer by segmenting confocal z-stacks starting at embryonic day (E) 13.5. We first measured basal cell size in three dimensions (Fig. 1a) using Fgf20-β-galactosidase knock-in allele embryos (Fgf20βGal), which report expression of Fgf20, one of the earliest known marker genes of placode cells^7^. Analysis of morphological features revealed that at E13.5 Fgf20βGal-positive cells were still cuboidal in shape and morphologically indistinguishable from other basal cells (Fig. 1b, c, Supplementary Fig. 1a, b). We also compared cell elongation in x-y, nematic order of elongation, as well as numbers of neighboring cells within Fgf20βGal-positive cells and the surrounding epidermis but also these tissue scale measurements revealed no specific morphological differences between Fgf20βGal-positive cell clusters and their neighbors (Supplementary Fig. 1a, b), indicating that placode cell fate specification precedes any substantial cell shape or tissue mechanical changes.

In stark contrast, the next embryonic day (E14.5) revealed a clearly distinguishable placode characterized by a decrease in the cross-sectional area and a corresponding elongation of the longitudinal (lateral) axis of basal cells (Fig. 1b, d). Interestingly, both apical and basal surfaces of the basal epidermal cells showed a proportional decrease from their length at E13.5, indicating that the elongation was not achieved through apical or basal constriction (Fig. 1b-e; Supplementary Fig. 1a, b). Further, 3D segmentation revealed that the elongation initiated between E13.5 and E14.5 occurred without alteration in cell volume. This analysis also confirmed that the placode cells displayed cylindrical shapes with comparable apical and basal surface areas (Fig. 1f, g). At E15.5, placode cells showed no further longitudinal elongation compared to E14.5 (Fig. 1b, d). However, while the apical surface length remained comparable to E14.5 cells, the basal surface expanded (Fig. 1d, e), resulting in a conical shape of E15.5 placode cells. Collectively these data indicated that initial placode cell fate determination precedes morphological changes. The subsequent morphological changes are initiated by a volume-preserving isotropic elongation of basal cells along the longitudinal axis at E14.5, followed by expansion of the basal surface of the basal cells at E15.5.

### In-plane oscillatory deformation drives tissue elongation

To understand the mechanics of the two-step morphological transformation, we investigated cell- and tissue-scale forces that could act on the placode cells. For this we performed live imaging of intact mouse embryos with genetically labeled plasma membrane-targeted Tomato (R26RmT/mG^16^). Visual inspection and particle image velocimetry (PIV) of live tissue dynamics to quantify cell movements indicated that cells within E14.5 placodes displayed coordinated collective centripetal oscillations, as well as rotational tissue flows around the placode as has been observed in skin explants previously^15^ (Fig. 2a, b; Supplementary Movie 1). Interestingly, flows were no longer detected at E15.5, whereas the oscillations continued and displayed overall negative divergence in the plane of the basal layer, indicative of tissue flows out of the basal cell layer plane (Fig. 2a, b; Supplementary Movie 2). Indeed, examining tissue flows in 3D revealed in-plane contractile oscillations positioned at the placode neck, extensile tissue flows downwards to the dermis, and deformation of junctions, resulting in plastic deformation of the placode structure and net downward tissue elongation (Fig. 2c; Supplementary Movie 3; Supplementary Fig. 2a; b). To understand the forces that could generate these patterns of tissue deformation, we measured mechanical stresses around the E14.5 placode using laser ablation^17^. Quantification of recoil and its radial displacement showed that recoil away from the cut occurred on both sides, confirming that the placode was under tensile strain (Fig. 2d, e; Supplementary Fig. 2c; Supplementary Movie 4). To identify the source(s) of tension on the placode, we examined patterns of actomyosin contractility. Phosphorylated myosin light chain-2 (pMLC2), which binds to the myosin-II heavy chain and generates contractile force, was higher along the apical domain than at the basal surface in both placodes and epidermal cells at E14.5, and additionally particularly enriched in the suprabasal cells (Fig. 2f, h). Surprisingly, despite the observed tension and contractile oscillations within the placode, pMLC2 was overall lower within the placode than in the surrounding epidermis (Fig. 2f-g). At E15.5, pMLC2 remained low within the placode and high within the suprabasal layers. In contrast, high myosin activity was detected within the dermis, particularly in structures that appeared as vascular networks as well as a defined ring-like structure around the placode (Supplementary Fig. 2d, e). Co-staining of vimentin and phalloidin confirmed this structure as a dense ring of fibroblasts that was detectable at E14.5 and became more prominent at E15.5 (Fig. 2i).

**Fig. 2.**
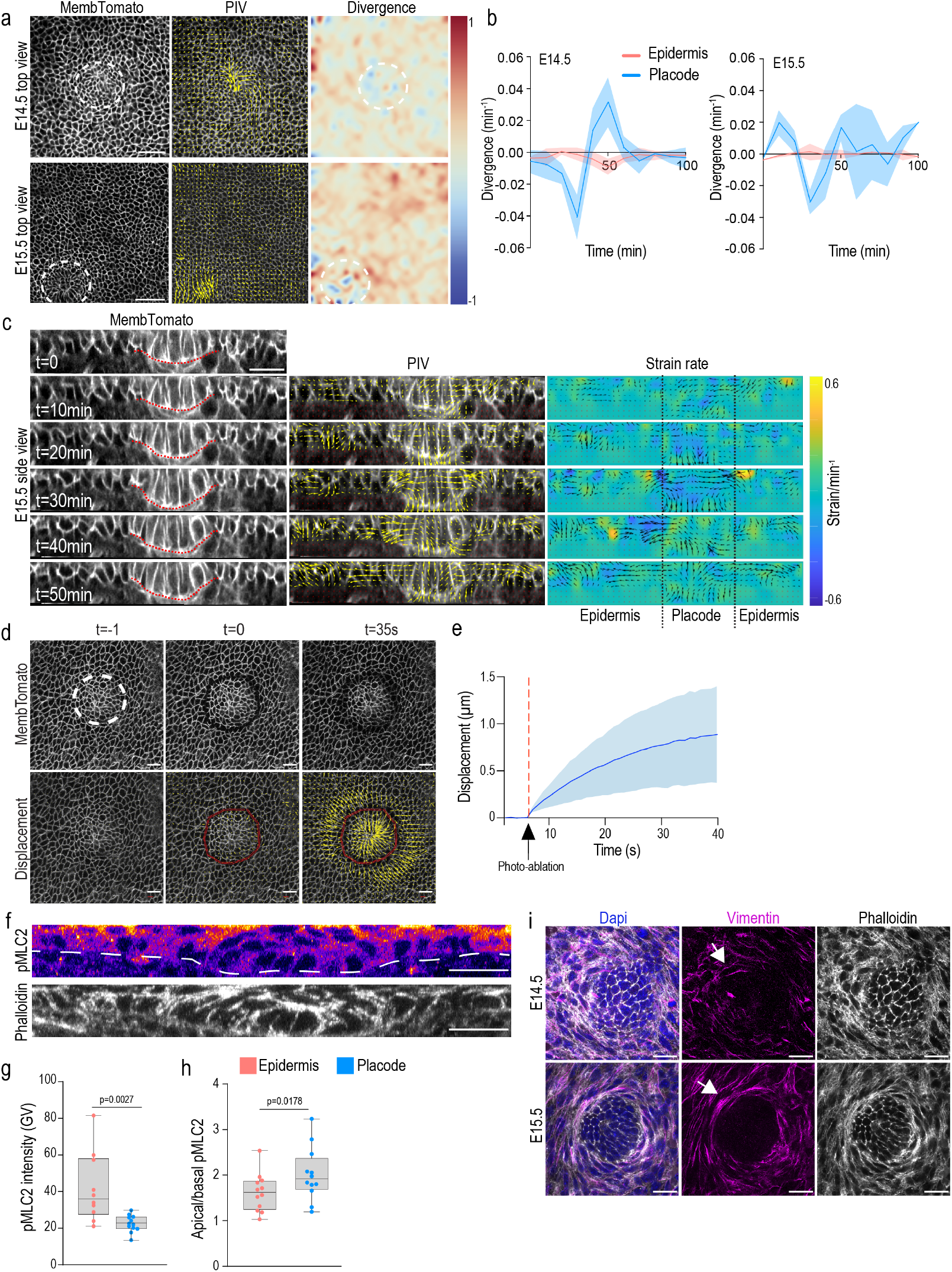
In-plane oscillatory deformation drives tissue elongation (a) Representative snapshots of whole embryo live imaging of membrane-labeled basal epidermis (left panels; white circles indicate placodes) that have been subjected to PIV analyses (middle panels) to extract mean divergence of flow vectors (right panels). Note localized tissue flows in both E14.5 and E15.5 placodes and negative divergence in E15.5 placodes (scale bars 50 μm n= 4 mice/stage). (b) Quantification of mean divergence over time (n=4 mice/stage; aligned to the first observed peak of negative divergence). Note oscillatory flows in both stages and net negative divergence in E15.5 placodes (n=4 (E14.5) and 5 (E15.5) placodes from 4 mice / stage). (c) Snapshots (left panels), and corresponding PIV (middle panels) and strain rate analyses (right panels) of optical cross sections from 3D time-lapse images of E15.5 whole embryos. Note progressive constriction of placode neck and associated downward flow of tissue indicative of plastic deformation (representative of 3 embryos; scale bars 20μm). (d) Representative snapshots of whole embryo live imaging of membrane-targeted Td-Tomato labeled basal epidermis with regions of laser ablation highlighted (white dotted circle; upper panel) and vectors showing recoil magnitude and direction after ablation (yellow arrows; lower panel). Scale bars 20 μm n= 4 mice. (e) Quantification of mean displacement over time (n=4 mice with 8-10 placodes/mouse). (f) Phalloidin and pMLC2-stained sagittal cross sections of mouse epidermis at E14.5. Scale bars 20 μm. Images representative of 3 mice/group. (g, h) Quantifications of pMLC2 intensity in basal placode and epidermal cells (f) and the apical/basal pMLC2 intensity ratio (g) in E14.5 epidermis. Note low pMLC2 within placode and high pMLC2 on apical/suprabasal surfaces (n=12 placodes from 3 mice). (i) Top view of skin whole mount at the level of the dermis shows a ring of vimentin- and actin-high fibroblasts (white arrows) around the placode at E14.5 and more prominently at E15.5. Images representative of 3 mice/group. Scale bars 20 μm.

Collectively these data confirmed that initial placode morphogenesis occurred in two stages: thickening/elongation at E14.5 was characterized by centripetal tissue oscillations, while at E15.5 the tissue underwent a transition towards downward tissue flows, constrained by in-plane contractions. Consistently, while contractility was low at lateral and basal cortical interfaces of placode cells, a ring of contractile fibroblast developed to tightly wrap around the placode base, coinciding with the centripetal oscillations in this area.

### Modeling suggests co-operative forces in cell and tissue shape transformations

To understand how the observed patterns of forces could generate the placode shape we turned to quantitative mechanical modeling. We developed a new, multi-layered 3D vertex model that consisted of basal epidermal and placode cells adhering to a basement membrane as well as multiple layers of suprabasal cells (see Methods for details and Supplementary Table 1 for model parameters). We used the model to probe how three experimentally observed morphological features – cell elongation, basal and apical placode surface area, and deformation/curvature of the epithelium (Fig. 3a) – are generated by the observed patterns of epidermal myosin activity, tissue tension, and fibroblast contractility. We explored two hypotheses in parallel: Model 1, where cell-intrinsic changes in lateral interfacial tension in placode cells drive the morphological changes, and Model 2, where extrinsic forces, such as those generated by the fibroblast ring around the placode, drive the changes (Fig. 3b). In both cases, we assumed that the observed asymmetry in myosin activity generates asymmetry in the tensions/wetting coefficients for the apical and basal surfaces of placode cells, which we describe in terms of a wetting coefficient at interfaces between placode and suprabasal cells (σ*_ps_*), and between placode cells and the basement membrane (σ*_pm_*); see Methods for more details). For Model 1, we additionally altered the wetting coefficient σ*_pp_* describing tensions at lateral interfaces between placode cells, while for Model 2, we simulated external contractile forces by imposing forces directed towards the center of the placode on all vertices of cells peripheral to the placode. The maximum force (f*_r_*) occurs at the edge of the placode, and forces decay exponentially towards zero over a length scale of several cell diameters away from the placode edge. To explore the effect of a wide range of parameters on the morphology of the placode, we generated phase diagrams (Fig. 3a-d; Supplementary Fig. 3a, b) by varying basal to apical wetting coefficient (σ *_pm_*/σ*_ps_*), lateral wetting coefficient (σ *_pp_*), and magnitude of extrinsic force (f*_r_*), over the entire physiological range (beyond this range the placode does not maintain integrity or produces abnormal curvatures). We then measured cell elongation, basal-to-apical surface ratio and curvature of the simulated placodes and compared them directly to the quantifications performed in embryos (Fig. 3c-e; Supplementary Fig. 3a, b). The simulations showed that both wetting coefficients and extrinsic force magnitudes had a noticeable impact on the extent of placode cell elongation, basal surface expansion and placode curvature. Notably, direct comparisons with the experimental data indicated that cell-autonomous differences in the lateral wetting coefficient were sufficient to recapitulate the morphological transformations observed at E14.5, but were insufficient to fully explain the subsequent morphological changes at E15.5, as indicated by the degree of overlap between measurements from the embryo and from the simulations (Fig. 3e). In contrast, co-operation of extrinsic forces in combination with a high basal to apical wetting coefficient recapitulated changes observed at both embryonic stages, (in Fig. 3e). Most importantly, only the extrinsic force Model 2 could robustly recapitulate the pattern of negative divergence of tissue flows observed in E15.5 embryos (Fig. 3f, Supplementary Fig. 3c). Collectively, the simulations predicted that cell autonomous changes in surface tension generated by polarized myosin distribution could be sufficient to generate some degree of cell elongation and curvature, but were not able to explain the morphological transformation at E15.5, where cell extrinsic forces might be playing a more important role.

**Fig. 3.**
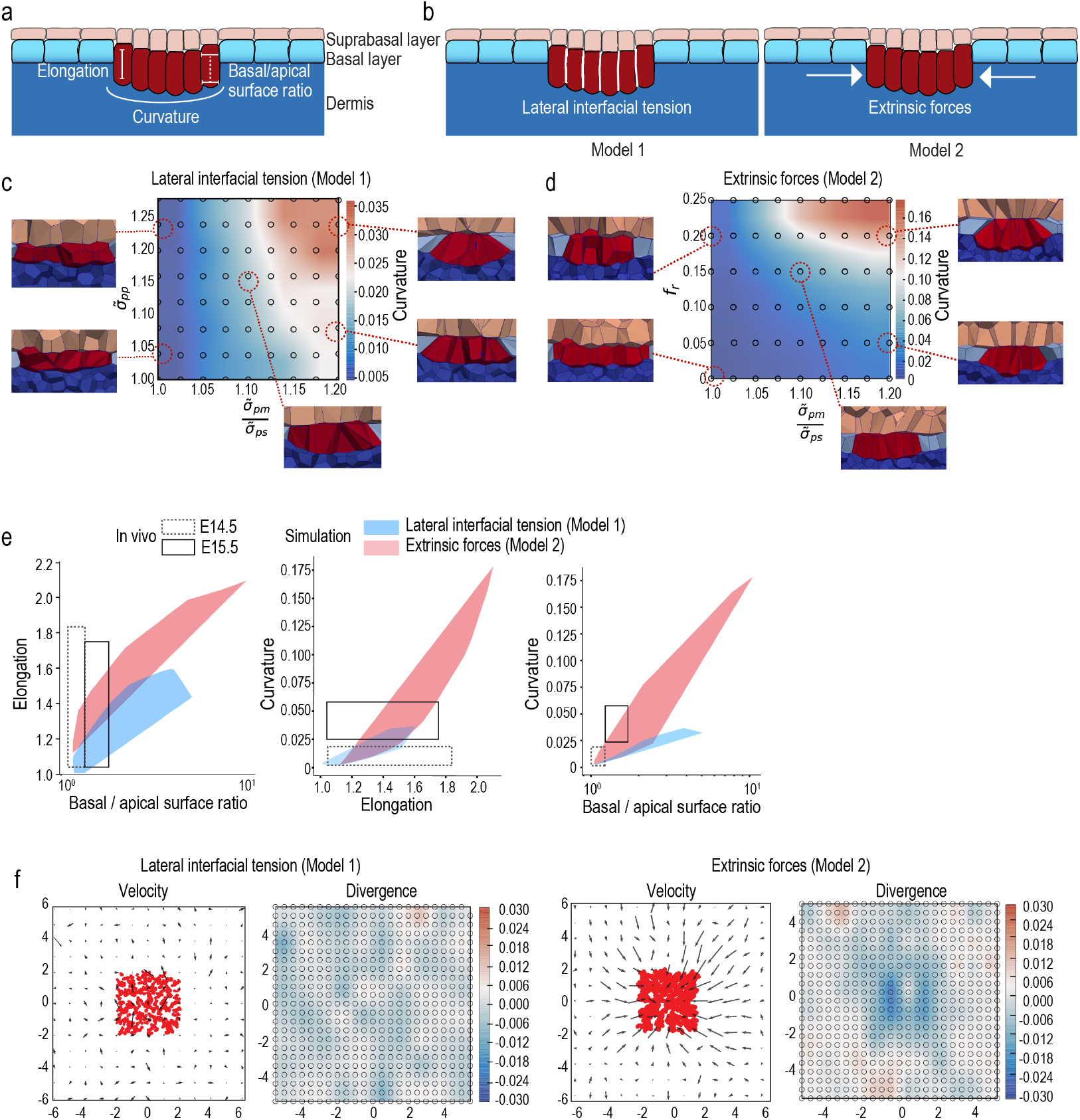
Modelling suggests co-operative forces in cell shape transformations (a) Schematics of the placode and parameters measured from embryos and simulations. (b) Schematic of the two models where either the lateral wetting coefficient (σ*_pp_;* Model 1) or magnitude of extrinsic forces (f*_r_*); Model 2) were varied in combination with a changing basal to apical wetting coefficient (σ*_pm_* /σ*_ps_*). (c, d) Phase diagrams of curvatures induced by interaction of lateral wetting coefficient (c) or extrinsic lateral forces (d) with varying changing basal to apical wetting coefficient. Note generation of tissue curvature resembling placode architecture with high lateral forces and high basal to apical wetting coefficient. (e) Comparison of morphological measurements from E14.5 and E15.5 embryos to simulations shows that model 2 with lateral cell non-autonomous forces recapitulates in vivo morphological transformations across both developmental stages whereas model 1 recapitulates cell shape changes at E14.5. (f) In-silico PIV analysis of tissue flow velocities and mean divergence of the flows from simulations with differential lateral wetting coefficient (model 1) or extrinsic lateral forces (model 2).

### Coordinated epidermal and dermal Myosin II activity is required for placode development and tissue compartmentalization

To challenge the model which predicted collaboration of cell-autonomous and extrinsic contractile forces in placode formation we proceeded to examine the role of myosin contractility in the epidermal and the dermal compartments. First, we deleted myosin-IIA, the major myosin isoform responsible for keratinocyte actomyosin contractility in the epidermis (Keratin-14 Cre deletion of Myh9 gene; Myh9-eKO^18^). At E14.5, although placode development was initiated, the placodes of Myh9-eKO mice were less invaginated compared to control mice (Fig. 4a, b; Supplementary Fig. 4a). Also the number of Sox9-positive cells and their local density were reduced within the Myh9-eKO placodes (Supplementary Fig. 4b-d). Interestingly, however, the cells still showed isotropic elongation comparable to cells in control mouse placodes (Fig. 4a, c). At E15.5, the placode showed a more pronounced phenotype as it failed to further invaginate. In addition, the dermal condensate, as marked by Sox2 expression^19,20^, was found partially embedded within the placode cells upon loss of epidermal contractility (Supplementary Fig. 4e, f). These data suggested that epidermal myosin IIA activity was required for the invagination and to generate forces against the dermal condensate, but it was not sufficient to induce full placode cell elongation and generate tissue curvature.

**Fig. 4.**
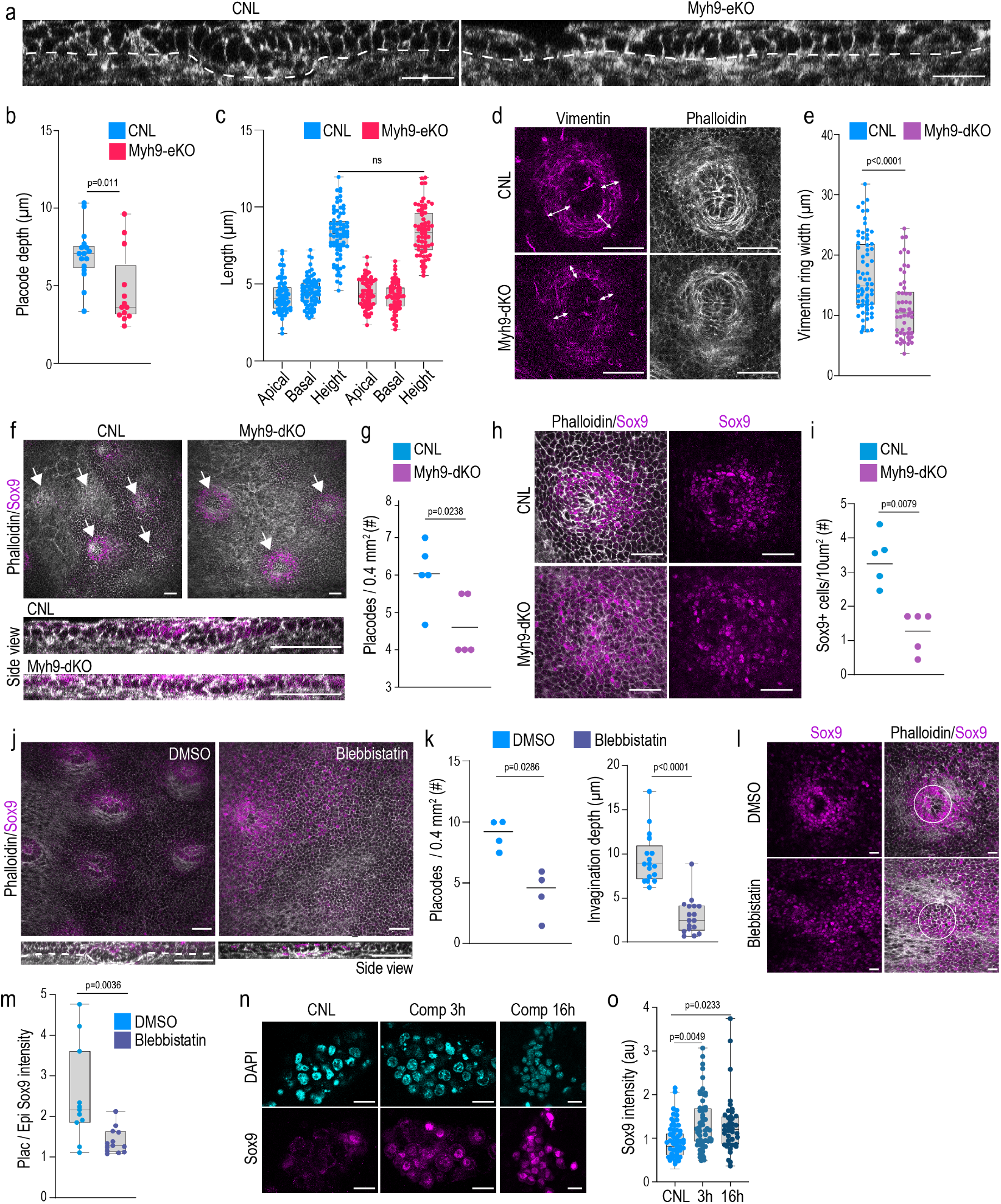
Coordinated epidermal and dermal Myosin II activity is required for placode development and tissue compartmentalization (a) Phalloidin-stained sagittal cross sections of control (CNL) and Myh9-eKO mouse epidermis at E14.5. Scale bars 20 μm. Images representative of 3 mice/group. (b, c) Quantifications of placode invagination depth (b) and placode cell surface lengths (c) from CNL and Myh9-eKO mice at E14.5 shows reduced placode depth but unchanged cell morphology in Myh9-eKO mice (n=16 (CNL) and 13 (Myh9eKO) placodes from 3 mice (left), 76 (CNL) and 66 (Myh9eKO) cells pooled across 3 mice (right); Mann-Whitney). (d, e) Representative vimentin- and phalloidin-stained dermal top views (d) and quantification (e) of skin whole-mounts from CNL and Myh9-dKO mouse epidermis at E16.5. Note reduced width of vimentin-positive fibroblast ring (arrows) in Myh9-dKO. Scale bars 50 μm (n=70 (CNL) and 55 (Myh9-dKO) placodes from 3 mice/group; Mann-Whitney). (f) Phalloidin and Sox9-stained whole-mounts of CNL and Myh9-dKO mouse epidermis at E14.5. Top views from basal layer (upper panels) and sagittal cross sections (lower panels) are shown. Scale bars 50 μm. Images representative of 5 mice/group. (g) Quantifications of placode numbers from CNL and Myh9-dKO mice at E14.5 shows reduced placode formation in Myh9-dKO mice (n=5 mice/genotype; Mann-Whitney). (h, i) Representative Sox9- and phalloidin-stained images (h) and quantification (i) of skin whole-mounts from CNL and Myh9-dKO mouse epidermis at E14.5. Reduced local density of Sox9+ cells in the placode is observed in the Myh9-dKO. Scale bars 50 μm (n=5 mice/group; Mann Whitney). (j, k) Representative phalloidin- and Sox9-stained images (j) and quantification (k) of skin explants treated with Blebbistatin (10 nM) for 24h starting at E13.5. Placodes were defined by tissue morphology and Sox9-positivity. Scale bars 50 μm (n=17 placodes from 4 mice; Mann-Whitney). (l, m) Representative phalloidin- and Sox9-stained images (l) and quantification (m) of skin explants treated with Blebbistatin for 24 h starting at E13.5. Circles mark placode area. Note reduced placode/epidermis Sox9 intensity ratio in the blebbistatin-treated explants. Scale bars 20 μm (n=11 (DMSO) and 13 (Blebbistatin) placodes from 3 mice; Mann-Whitney). (n, o) Representative images (n) and quantification (o) of Sox9-stained control and compressed organoids. Scale bars 20 μm (n=50 (CNL), 56 (3h) and 40 (16h) organoids pooled across 4 independent experiments; Kruskal-Wallis/Dunn’s).

As the simulations predicted a critical role for extrinsic forces in placode formation, we proceeded to examine the role of the contractile fibroblasts around the developing placode. To this end we deleted Myh9 specifically in fibroblasts using Twist2-promoter driven Cre^21^ (Myh9-dKO; targeting dermal fibroblasts lining the epidermis and placode high in Twist2 expression; Supplementary Fig. 5a). Analyses of the fibroblast ring showed reduced thickness in Myh9-dKO mice dermis (Fig. 4d, e), confirming the hypothesis that myosin-IIA activity is required to assemble the ring of fibroblasts surrounding the placode. Importantly, much fewer placodes were observed in the Myh9-dKO epidermis at E14.5 (Fig. 4f, g), and cells within these Myh9-dKO placodes were slightly less vertically elongated (Supplementary Fig. 5b). A severe defect in early placode morphogenesis was confirmed by the substantial attenuation of the second morphogenetic wave at E16.5, as indicated by reduced placode number and increased distance between individual placodes (Supplementary Fig. 5c, d). Where placodes had formed, they displayed slightly reduced invagination (Fig. 4f; Supplementary Fig. 5e). This indicated that myosin IIA activity in the dermal fibroblasts and extrinsic mechanical forces are essential to promote placode formation, as also predicted by the simulations. Surprisingly, Myh9-dKO placodes showed an overall more dispersed localization within the epidermis. Further quantification revealed significant reduction in number and density of Sox9-expressing cells in particular around the prospective placode area defined by the presence of the dermal condensate and epithelial thickening (Fig. 4h, i; Supplementary Fig. 5f). Additionally, and in contrast to the Myh9-eKO, a significant increase of the average distance of Sox9-positive cells from the center of the placode and in the total area of tissue containing Sox9-positive cells was observed in the Myh9-dKO placodes, indicative of a failure of Sox9 compartmentalization (Supplementary Fig. 5f, g). To investigate the potential collaborative role of myosin contractility across the compartments, we prepared full skin explants from E13.5 and E14.5 mice and treated them with Blebbistatin for 24h to block myosin activity in all cells. Strikingly, when myosin was inhibited in all cells at E13.5, the placodes, defined by cell shape changes associated with Sox9 expression, almost completely failed to develop (Fig. 4j, k), indicating a critical role of collective myosin activity in both compartments in placode thickening as also predicted by the model. In addition to the morphogenetic defects, Sox9 expression failed to become compartmentalized into placode cells (Fig, 4l, m). Treating E14.5 embryos instead resulted in an impairment of placode invagination into the dermis, with the dermal condensate partially indenting the epithelial structure (Supplementary Fig. 5h, i), indicating that myosin II activity is further required for placode invagination and curvature.

As reducing extrinsic forces on the placode by myosin IIA inhibition resulted not only in reduced placode formation but also in reduced Sox9-levels and compartmentalization, we hypothesized that mechanical confinement of the placode cells by the contractile fibroblast ring promotes Sox9 expression and compartmentalization. To test this, we isolated epidermal progenitor cells from E14.5 embryos and placed them in 3D collagen hydrogels and subjected them to hydrostatic pressure through extrinsic static compression (Supplementary Fig. 5j). As expected, very little Sox9 expression was observed in the control cell clusters (Fig. 4n, o). In contrast, upon application of external static compression on the hydrogels, robust Sox9 expression was observed already after 3h of compression, and the effect was sustained after 16h of compression (Fig. 4n, o). Collectively these data, in line with the model predictions, indicated that changes in epidermal interfacial tension through altered epidermal contractility are required to produce contractile force for the placode to invaginate and to push down the dermal condensate. However, the full morphogenetic transformation of cell elongation and tissue deformation require additional extrinsic forces generated by a contractile fibroblast ring wrapping the placode. Surprisingly, these extrinsic forces are also essential for promoting compartmentalization of the tissue by directing Sox9 expression.

### Spatiotemporal coordination of cell divisions control placode downgrowth

We next asked what facilitated the profound transition of the placode from an initial epithelial thickening at E14.5 to a downward flowing, budded structure at E15.5. To this end we investigated patterns of cell divisions that have been shown to promote tissue fluidification as well as Drosophila tracheal budding and epithelial buckling^22–25^. Using both embryo live imaging of Histone2B-mCherry/membrane-EGFP mice to visualize dividing cells (R26R-RG mice^26^) and quantitative imaging of Ki67-stained fixed tissues, we examined patterns of proliferation at E14.5 and E15.5. These analyses showed that cell divisions were substantially more frequent within the epidermis at E14.5, compared to the placode (Fig. 5a, c; Supplementary Movie 5). This low mitotic frequency in the placode was associated with decreased levels of active nuclear YAP (Fig. 5a, d), the mechanosensitive transcription factor that acts within the Hippo pathway to control growth and tissue size^27,28^. In stark contrast, at E15.5, the most invaginated lower surfaces of the placodes showed higher levels of cell division (Fig. 5b, c; Supplementary Movie 6), accompanied by restoration of nuclear active YAP (Fig. b, d), consistent with recent work showing cell compression promotes nuclear exit of YAP to inhibit cell division^29^. In line with this, inhibition of YAP in freshly isolated epidermal progenitors showed decreased levels of cell division (Supplementary Fig. 6a, b). Collectively, this indicated that mechanical confinement of E14.5 placode cells repressed cell divisions through nuclear exclusion of YAP.

**Fig. 5.**
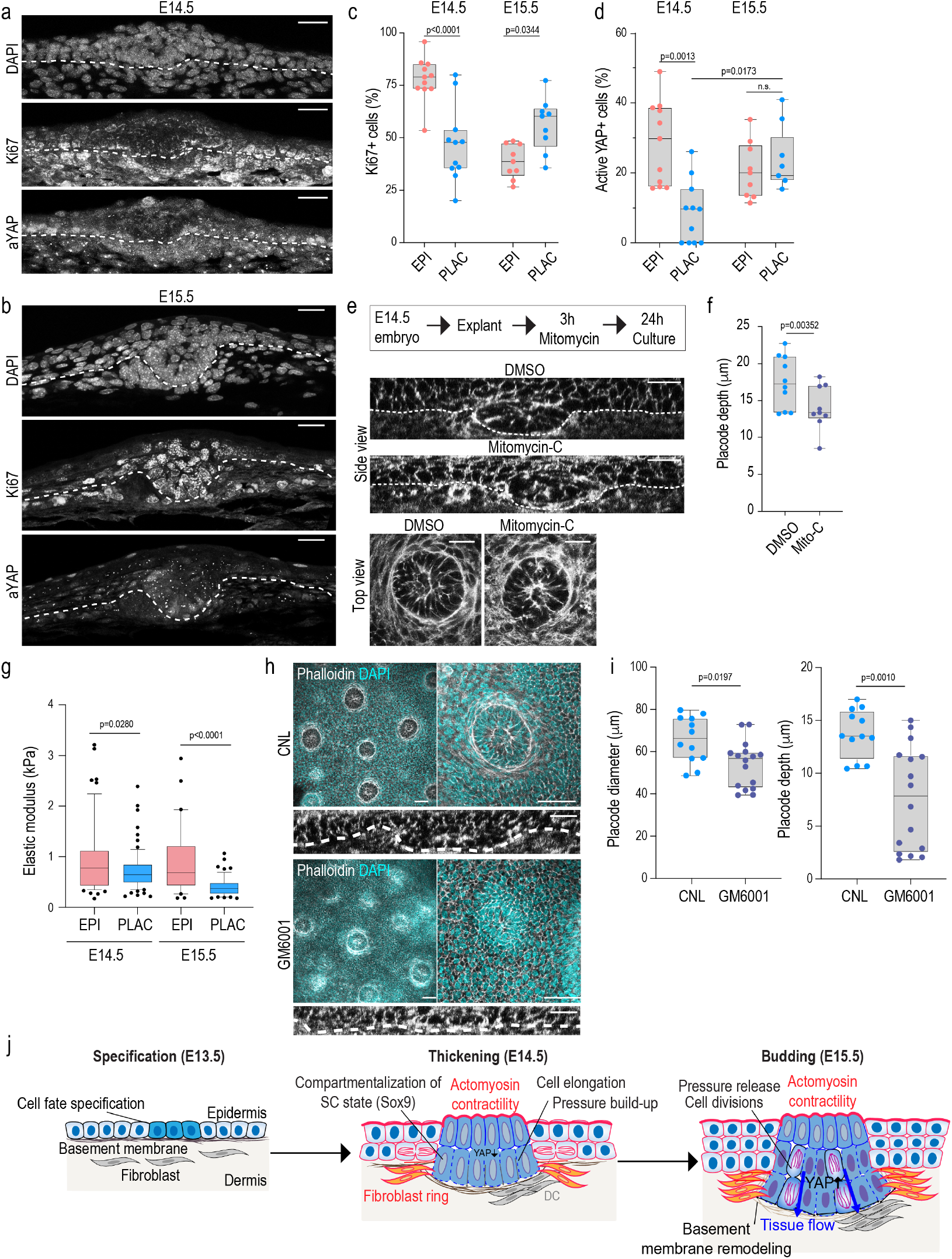
Spatiotemporal coordination of cell divisions control placode downgrowth (a, b) DAPI, Ki67 and active YAP (aYAP)-stained sagittal cross sections of E14.5 (a) and E15.5 (b) mouse epidermis. Scale bars 20 μm. Images representative of 3 mice/group. (c, d) Quantification of Ki67 (c) and aYAP (d) from E14.5 and E15.5 placodes and epidermis. Note low levels of Ki67 and aYAP in E14.5 placode and elevation of these intensities to comparable levels with epidermis in E15.5 placodes (n=11 (E14.5) and 9 (E15.5) placodes or surrounding epidermis from 3 mice/group; one-way ANOVA (c) and Kruskal-Wallis (d)). (e) Experimental outline and optical cross sections and top views of E14.5 phalloidin-stained Mitomycin-C-treated and DMSO control skin explant cultures. Scale bars 20 μm. Images representative of 3 explants/group. (f) Quantification of placode diameter and depth from experiments shown in (h) (n=10 (CNL) and 9 (Mito-C) placodes from 3 explants; Student’s t-tests). Note reduced placode depth Mitomycin-treated skin explants both at E14.5 and E15.5 and decreased in placode diameter at E14.5 in Mitomycin-C-treated explants. (g) Quantification of elastic moduli of basement membranes using force indentation spectroscopy. Note reduced elastic moduli of basement membranes below placodes (n>50 force curves pooled across 3 mice/group; Kolmogorov-Smirnov test). (h) Optical cross sections and top views of DAPI and phalloidin-stained skin explant cultures isolated from E14.5 mice and treated with GM6001 (10μM) or DMSO control for 48h. Scale bars 50 μm (top view), 20 μm (side view). Images representative of 3 explants/group. (i) Quantification of images shown in (e) show reduced placode diameter and depth in GM6001-treated explants (n=12 (CNL) and 16 (GM6001) placodes from 3 explants/condition; Mann-Whitney). (j) Model of placode formation indicates early cell fate specification (E13.5) followed by polarized myosin activity and assembly of peri-placode contractile fibroblast ring to generate tissue elongation and basal cell surface expansion (E14.5-E15.5). Tissue elongation builds up pressure that is released by basement membrane remodeling, allowing YAP activation, localized cell divisions and placode budding (E15.5).

The enrichment of cell divisions outside of the placode at E14.5 and subsequently broadly within the basal cell layer of the placode at E15.5 led us to hypothesize that high levels of divisions at the placode base at E15.5 could accelerate budding. To test this hypothesis, we prepared skin explants from E14.5 mice and treated them with Mitomycin-C for 3h to halt cell divisions, followed by 24 h of culture to allow further morphogenesis (Fig 5e; Supplementary Fig. c, d). Indeed, blocking cell divisions at E14.5 did not substantially affect placode diameter, but strongly attenuated budding (Fig. 5e, f; Supplementary Fig. 6e). Collectively, these data indicated that confinement of the placode at E14.5 promotes exclusion of YAP and prevents cell divisions, while the subsequent folding/buckling of the epithelium at E15.5 releases compressive stress on the placode, facilitating YAP activation, cell cycle re-entry and mitoses, which then trigger efficient budding of the placode.

### Basement membrane remodeling facilitates placode budding

Finally, we asked what facilitated release of mechanical confinement allowing cell cycle re-entry and downward budding of the placode. Previous work using oncogenic mutations to model epidermal tumors showed that active basement membrane remodeling, triggering its local soft-ening, drove formation of tumor buds^30^. To test if placode budding was associated with basement membrane softening, we first analyzed basement membrane composition using im-munofluorescence, but did not observe substantial changes in the presence of the two major laminins, LN-511 and LN-332, or Collagen IV, at the sites of placode compared to the surrounding epidermis, and the basement membrane ap-peared continuous across the placode (Supplementary Fig. 6f, g). However, measurements of the stiffness of the base-ment membrane using atomic force microscopy revealed a slightly softer basement membrane beneath the placode at E14.5. This local softening became significantly more pro-nounced at 15.5 (Fig. 5g). To address the mechanism of local basement membrane softening in the absence of substantial changes in molecular com-position, we asked if proteolysis-driven turnover could play a role. We first examined the expression of extracellular ma-trix (ECM)-degrading enzymes by re-analyzing single cell sequencing data from the E14.5 mouse skin^31^. We analyzed protease expression in the various skin cell populations and found the key proteolytic enzyme Matrix Metalloproteinase (MMP)-2 highly expressed by the dermal fibroblasts, and the membrane bound MMP-14 specifically enriched in the placode as well as in the dermal condensate (Supplementary Fig. 6h). Other MMPs were not expressed at high levels. To test if MMP-mediated matrix remode-ling was required for placode invagination we treated E13.5 skin explants for 48h with the broad-spectrum MMP inhibitor GM6001 to block pro-teolytic degradation of the ECM and basement membrane. Strikingly, inhibiting matrix remodeling strongly attenuated placode budding, as visualized by both the decreased diame-ter and depth of the developing placodes within the explants (Fig. 5h, i). Collectively these data indicated that ECM re-modeling triggered by proteolytic degradation is essential for placode growth by allowing pressure release and subsequent downgrowth.

## Discussion

By combining 3D vertex modeling with ex vivo biophysical measurements and genetic manipulations, we have dissected a two-step mechanism of placode morphogenesis and compartmentalization that depends on coordination of mechanical forces across compartments. In the first step, polarized myosin activity in the epithelium together with contractile activity from fibroblasts generate oscillatory contractions to produce a local, mechanically confined tissue thickening (Fig. 5j). In the second step, release of pressure through local extracellular matrix remodeling facilitates tissue fluidification, which due to the continuing oscillatory contraction produce a plastic deformation and downward budding of the tissue (Fig. 5j). The precisely-timed interactions between cell types and the forces they generate, and the transmission of these forces through the extracellular matrix and tissue mechanical changes, highlights the complexity of orchestrating morphogenesis and cell state transitions in multicompartment mammalian tissues. Interestingly, in vitro studies in experimentally confined epithelial cell lines have revealed similar interplay between pressure, growth control, and buckling^29,32^, suggesting that the mechanisms described here could be applicable to developmental processes in other epithelial organs. Further, the effect of confinement on stem cell differentiation is consistent with previous work linking tension to cell fate progression through changes in chromatin organization^18,33^.

Our findings are further interesting in light of recent reports showing that mesenchymal dynamics regulate feather follicle patterning in avian skin^34,35^. Collectively these studies highlight how mechanical crosstalk across compartments is essential to propagate symmetry breaking and the emergence of tissue patterns. Our work also identifies an interesting distinction where the murine placode early fate specification seems to precede mechanical changes, which subsequently are essential for further cell type specification and in particular compartmentalization of cell fates. Interestingly, the distinct role of fibroblasts in generating forces, and adjusting pressure through remodeling the ECM recapitulate features observed between cancer-associated fibroblasts and tumor cells during cancer progression^30,36^. These common features between placode development/budding and cancer progression, are consistent with observations on cancer cells hijacking embryonal pathways to promote aggression^37–39^. This suggest that the mechanisms discovered in this study as well as the modeling approaches developed could be of broad relevance for various biological processes. In particular it will be interesting to understand how stromal fibroblasts and changing patterns of epithelial contractility, pressure and tissue fluidity play roles in the initiation of cancer aggression.

## Methods

### Mice

Myh9 floxed mice were obtained from the European Mouse Mutant Archive (EM:02572)^40^ and crossed with the K14-Cre line^18,41^ or the Twist2-Cre line obtained from JAX laboratories (stock 008712)^21^. Membrane-targeted Tomato reporter mice (R26RmT/mG 16 were from JAX laboratories (stock 007676) 16. Histone2B - mCherry/membrane-EGFP (R26R-RG mice^26^; were from Riken Laboratories for Animal Resources and Genetic Engineering (LARGE) and were crossed with K14-Cre line to obtain epidermis-specific expression. FGF20-LacZ reporter mice have been previously described^7^. Embryonic ages were defined based on the presence of vaginal plug (taken as a embryonic day 0), limb morphology and other external criteria^42^. All mouse studies were approved and carried out in accordance with the guidelines of the Finnish national animal experimentation board (ELLA) or the Ministry for Environment, Agriculture, Conservation and Consumer Protection of the State of North Rhine-Westphalia (LANUW), Germany.

### Immunofluorescence and confocal microscopy

For skin whole mounts, embryos were dissected from pregnant mice and directly fixed in 37°C pre-warmed 4% paraformaldehyde (PFA) at room temperature for 3 h. After multiple PBS washes, the skin was carefully peeled using forceps and placed in blocking solution (0.5% Triton-X, 5% bovine serum albumin (BSA), 3% normal goat serum (NGS)) for 1 h at room temperature (RT). For paraffin sections tissue biopsies were fixed in 4% PFA, embedded in paraffin and sectioned. Sections were de-paraffinized using a graded alcohol series and antigen retrieval was carried out using target retrieval solution (Dako) at pH 6 or pH 9 in a pressure cooker. For cryo-sections, tissue samples were placed unfixed into cryomolds containing OCT Tissue Tek and allowed to freeze on dry ice. 6-8 μm thick sections were cut, airdried and fixed with 4% PFA. For basement membrane stainings, unfixed cryosections were used. For both paraffin and cryofixed sections, samples were blocked in 5% BSA, 3% NGS. Organoids and 2D cell culture of primary epidermal stem cells were fixed in 4% PFA for one hour at RT. After PBS washes, samples were placed in blocking solution (0.5% Triton-X, 5% BSA, 3% NGS) for one hour at RT. Samples were incubated with primary antibodies diluted in blocking solution overnight in a moist chamber at 4°C (organoids and 2D cell culture) or at room temperature (whole mounts). For paraffin sections antibodies were diluted in Dako antibody diluent (Agilent) overnight in moist chamber at 4°C. The following primary antibodies were used: rabbit anti-p-MLC2 (p-Ser20; Fisher Scientific, NC0279729; 1:100) rabbit anti-Vimentin (Santa Cruz, sc-7557-R; 1:100), rabbit anti-Sox9 (Cell Signalling, 82630; 1:100), rabbit anti-Sox2 (Millipore Sigma, AB5603; 1:200), mouse anti-Twist (abcam, 50887; 1:200), mouse anti-Ki67 (Cell Signalling, 9449; 1:300), rabbit anti-active YAP1 (Abcam, 205270; 1:100), guinea-pig anti-Keratin 14 (Progen, GP-CK14; 1:300), rabbit anti-Collagen IV (Abcam, 6586; 1:200), rabbit anti-Laminin 332 (gift from R.E Burgeson; ^43^; 1:20 000), rat anti-Laminin a5 (504^44^; gift from L. Sorokin; 1:20 000), chicken anti-beta-galactosidase (Abcam, 9361; 1:500). F-actin was detected using Alexa647-conjugated phalloidin (1:500). Bound primary antibody was detected by incubation with Alexa-Fluor 488-, 568- or 647-conjugated antibodies (Invitrogen). Nuclei were counterstained with 4, 6-diamidino-2-phenylindole (DAPI, Invitrogen). After PBS washing, slides were mounted in Elvanol. All fluorescence images were collected by laser scanning confocal microscopy (SP8X; Leica) with Leica Application Suite software (LAS X version 2.0.0.14332), using a 20X dry objective or a 63x water immersion objective. Images were acquired at room temperature using sequential scanning of frames of 1 μm thick confocal planes (pinhole 1). Images were collected with the same settings for all samples within an experiment.

### Ex vivo embryo imaging

Imaging was performed essentially as described earlier^33^, with minor modifications. Briefly, embryos were dissected at embryonic stages E14.5 or E15.5 and immobilized on imaging chambers engineered on top of Lumox-teflon imaging plates that allow gas exchange (Sarstedt). Whole embryos were immersed in epidermal growth medium (CnT-PR, CELLnTEC Advanced Cell Systems) supplemented with calcium-depleted 10% fetal bovine serum (HyClone/Thermo Fisher), 20 U/ml penicillinstreptomycin (Thermo Fisher), 20 mM Hepes, 1.8 mM CaCl2, and 10 ng/ml mouse recombinant EGF (Novus; NBP2-34952). Imaging was performed using an Andor Dragonfly 505 high-speed spinning disk confocal microscope (Oxford Instruments) with an inverted Nikon Eclipse Ti2 microscope (Nikon) equipped with 488nm and 546 nm lasers, an Andor Zyla 4.2 sCMOS camera and an environmental chamber set at 37°C, 5% CO_2_. Acquisition was carried out with 25X dry or 40X water immersion objectives using the Fusion 2.0 software. After acquisition, the image series were 3D drift corrected using manual registration on Image J together with the descriptor-based series registration ImageJ plug-in before analysis. Image analyses are described in detail below. Optical cross section live image sequences were generated using the oblique slicer tool on Imaris.

### Skin explants

Embryonic back skins were dissected from freshly collected E13.5 or E14.5 embryos, placed at 37 °C, 5% CO_2_ on a 3 μm pore size cell culture insert (Corning) in DMEM supplemented with Glutamax, 10 percent FBS, 1% penicillin-streptomycin at an air-liquid interface as previously described^45^. Skins were dissected into two pieces and one half was treated with DMSO as control and the other half with Mitomycin C (1μg/ml; Sigma), Blebbistatin (10nM; Sigma) or GM6001 (10μM; Sigma) for 3, 24, or 48 h) as indicated for each specific experiment. After culture, tissues were fixed and processed for immunofluorescence as described above for tissue whole mounts.

### Epidermis cell culture and hydrogels

For cell culture experiments, mouse epidermal stem cells were isolated from E15.5 embryos (organoids) or P0 mice (2D culture). After dissection, skins were treated with Antibiotic/Antimycotic solution (Sigma A5955) for 5 min and epidermal single cell suspensions were generated by incubating skin pieces in 0.8% trypsin for 30 min at 37°C. Epidermal single cell suspensions were isolated and cultured in 3C organoid medium as described earlier^46^. For 3D organoids, cells were suspended in growth factor-reduced Matrigel (Corning) and grown in in 3C medium (MEM Spinner’s modification (Sigma), 5 μg/mL insulin (Sigma), 10 ng/mL EGF (Sigma), 10 μg/mL transferrin (Sigma), 10 μM phosphoethanolamine (Sigma), 10 μM ethanolamine (Sigma), 0.36 μg/mL hydrocortisone (Calbiochem), 2mM glutamine (Gibco), 100 U/mL penicillin and 100 μg/mL streptomycin (Gibco), and 10% chelated fetal calf serum (Gibco), 5 μM Y27632, 20 ng/mL mouse recombinant VEGF, 20 ng/mL human recombinant FGF-2 (all from Miltenyi Biotec)). For 2D cultures, cells were seeded on glass-coverslips previously coated with 30μg/mL collagen I (Millipore) and 10μg/mL fibronectin (Millipore) for 1 hour at 37°C. Cells were cultured in 3C medium, incubated with DMSO or Verteporfin (10μM; Stem Cell technologies) for 6 hours before fixation. For compression studies, silicon cylinders with 500 g weight were placed on top of hydrogels at 37°C, 5% CO_2_ for 3 h or 16 h. After compression, samples were directly fixed and immunofluorescence were performed as described above.

### Laser ablation on ex vivo whole embryos

E14.5 embryos were collected and immobilized on imaging chambers as described above. Immediately after the collection, imaging and photo-ablation were performed using a Zeiss LSM880 confocal microscope equipped with a 561nm laser, a two-photon laser Ti:Sapphire laser (Mai Tai DeepSee, Spectra Physics) and an incubation system set at 37°C, 5 % CO_2_. For photo-ablation, the two-photon laser was set at 800 nm and 75% laser power. Images were acquired with a 40x/1.30 Oil objective. The region to be ablated was drawn as a circle surrounding the placode. The thickness of the line remained constant for all ablations. Images were captured at 1 second intervals for 40 seconds, starting 5 seconds before ablation. Quantification of displacements after ablation was performed as previously described^32^. Briefly, Particle-image velocimetry (PIV) was first performed on the entire field of view using a window size of 64*64 pixels and an overlap of 0.75. PIV was performed by comparing each timepoint to the last timepoint before ablation to obtain tissue displacement. To perform radial displacement profiles, the local normal direction respect to the ablated region was computed as described ^32^. This normal direction was used to calculate the radial component of the displacement, which was then averaged as a function of distance to the ablated region. The radial displacement profile was used to calculate the total tissue displacement as the time evolution of the distance displaced by the tissue at both sides of the cut. For that, the inner peak of negative displacement was subtracted from the outer peak of positive displacement.

### Image analyses

For tissue-scale analysis, 2D segmentation and morphometry were performed using Tissue Analyzer (v2.3)^47^. Placode morphogenesis and cell shape (apical/basal lengths; z axis length) were quantified manually using Fiji^48^. Hair follicle placodes, and dermal condensates were identified based on the expression of specific markers (FGF20, Sox9 and Sox2) and morphology. To quantify placode depth, the distance between epidermal basement membrane and deepest point of placode cells was measured. The closest distance between placodes was measured manually from the center of one placode to the center of the next closest placode. To measure pMLC2 intensity, regions of interest restricted to the placode cells and the surrounding interfollicular epidermal cells were determined and mean fluorescence intensities were quantified. Each placode was paired to its surrounding epidermis. To quantify the ratio of apical/basal PMLC2 mean intensity, regions of interest corresponding the apical and basal surfaces were generated and the mean fluorescence intensities quantified. Sox9-positive cells were quantified manually. The coordinates of Sox9-positive cells were extracted using Fiji. To quantify the pattern of Sox9 expression, four different parameters were measured using custom-built Python scripts: 1) The total number of Sox9 positive cells in placode area (defined by the presence of a dermal condensate underneath the epithelium and a thickening of the epithelium itself), 2) local density of Sox9-positive cells within a circular area with a radius of 10 μm around the placode (defined as above); 3) the average distance of Sox9-positive cells from the center of the placode (determined as above); 4) the convex Hull area corresponding to the minimal area necessary to include all the Sox9-positive cells. To measure Sox9 mean intensities in organoids, a nuclear region of interest was generated using automated thresholding of the DAPI staining, after which mean fluorescence intensities of Sox9 were quantified within the nuclear area of interest from maximum projection images of 15 stacks. Sox9 mean nuclear intensities were normalized to DAPI intensity of the corresponding individual nucleus. Images were collected with the same settings for all samples within an experiment Ki67 and YAP1-positive cells were determined manually from paraffin tissue sections and normalized to total number of nuclei obtained from DAPI masks. For Ki67 and aYAP1 intensities from primary epidermal stem cells, 5 confocal planes (equivalent to 5 μm) were projected using maximum intensity projection type. The number of mitosis (from metaphase to telophase) from DMSO- or Mitomycin-treated skin explants were identified using DAPI staining and manually quantified inside the placode and in the surrounding interfollicular epidermis for each field of view (0.4 μm^2^).

### 3D tissue rendering and 3D cell shape

Images were acquired from freshly isolated whole unfixed mTmG E14.5 embryos with the Andor Dragonfly 505 high-speed spinning disk confocal microscope. Images were acquired as z-stacks of 1 μm thick optical sections and cells were segmented and tracked across every z-stack using Tissue Analyzer, after which x-y cell dimensions were measured and treated as pseudo-time. To ensure accurate segmentation of cell heigh, a nuclear bounding box was determined in parallel from skin whole mount stained with DAPI and imaged with a higher resolution using the laser scanning confocal microscopy (SP8X; Leica) with 63X glycerol immersion objective. 3D rendering was then performed using a custom-built code in Mathematica as follows: The cell junctions were manually corrected with a combination of Tissue Analyzer and Mathematica (v12.3) and tracking were performed to stitch the cells across the stacks. Cells that only appeared in less than 3 frames were discarded from the analysis. The cellular regions of interest were then converted from image objects to mesh regions in Mathematica by first applying a 3D Gaussian Filter over a radius of two pixels to smooth the roughness and then applying a “Dual Marching Cubes” algorithm using “ImageMesh” function. The cellular mesh regions were then rendered appropriately in a box with the scaling parameters of the image stack. The computations for cellular volume and associated parameters were performed on the mesh objects and on point clouds derived from the mesh.

### Particle Image Velocimetry (PIV)

PIV was performed using PIVLab version 2.59. Live imaging movies of embryos were first imported into PIVLab, where a multiple pass windowed FFT based approach was utilized to track tissue scale flows for both sagittal and axial cross-sections. In order to correct for variations in intensities and prevent tracking errors a CLAHE (Contrast Limited Adaptive Histogram Equilisation) was performed with a box of size 10 pixels. In addition, a denoising was performed on the resulting images using Weiner Noise Filter with a max pixel count of 3. The tracking was initially done on test images by varying window sizes of the first and second pass, and where applicable the third pass. Below a certain threshold performing PIV resulted in patterns of vector fields that were determined to be insensitive to the choice of window size. Based on this, a constant size of 100 pixels was selected for the first pass, with the second pass set as 50 pixels with an overlap between first pass windows set to 50 pixels. For PIV on frontal cross-sections from time-lapse images, masks were manually drawn over regions the epithelium and a three-pass windowed FFT was performed with the first window size set to the maximum allowable value determined by the algorithm. The resulting vector fields were visually inspected, and any spurious vectors were manually rejected from the analysis. Since the maximum allowable window-size for sagittal cross section was relatively smaller than the spacing between lateral junctions, the strength of the vector field was smoothed within PIVLab to account for the noise. The vector fields were utilized to compute divergence maps and strain-rates. To compute mean divergence within the placode a circular ROI covering the placode was drawn over the raw image and imported into the PIVLab where the mask was overlayed on the divergence field. Likewise, for computing mean divergence of the interfollicular epidermis, placodes were masked out to perform PIV on all the region except the placodes. The values were subsequently integrated within the regions to yield a mean value. An optic flow analysis was also performed in parallel and the results were compared with PIV to test for consistency. To plot mean divergences from multiple embryos the divergence of each placode and its surrounding IFE were aligned according to the first observed peak of negative divergence.

### 3D vertex model and simulations

The 3D vertex model code was adapted from work by Zhang and Schwarz^49^ based on previous work^50,51^. A multilayered 3D vertex model tissue structure was generated to capture the interactions between the basement membrane, basal cells and suprabasal cells. A subset of the basal cells is labeled as placode cells, with potentially distinct mechanical properties. Each cell in the network is represented as a polyhedron with shared vertices and edges such that the tissue is completely confluent, with no gaps or holes in the tissue. Although the basement membrane is not composed of biological cells, we use vertex cells with a special energy functional, described below, to represent its mechanics. The multilayered structure is initialized from a Voronoi tessellation of a uniform 3D point pattern using the Voro++ library^52^, and we minimized the vertex model energy function before the various cell types are specified. Periodic boundary conditions are maintained at the edges of the simulation domain in all directions. Neighbor exchanges (which are called T1 transitions in 2D but are generally more complicated in 3D) are allowed via I-H and H-I transitions as described^50^. Each cell is labeled as one of four: suprabasal, basal, placode, and basement membrane.

### Computing curvature in tissue and experiments and cell elongation in simulations

The curvature of the simulated tissue is determined in a two-step process. A paraboloid is first fitted to the basal vertices shared between the placode cells and those representing the basement membrane. Subsequently a Gaussian Curvature is determined at the deepest point for the fitted surface. In the embryonic tissue whole mounts, we manually identify points on the interface between the placode cells and the membrane in 2D cross section views. These points are fitted to a parabola assuming that the associated parabola is symmetric along the missing axis and generate the associated paraboloid. The same procedure as with simulation data is then followed to extract curvature. In order to quantify cell elongation, the depth of the paraboloid is extracted and combined with the average height of cells in the basal layer.

### Single cell RNA sequencing analysis

Raw sequence data for E14.5 mouse skin^31^ was obtained from NCBI GEO (accession ID: GSE122043). The quality of the data was checked using FastQC (v0.11.9). STARsolo aligner (v2.7.7a)^53^ was used to align sequenced reads to the mouse reference genome (GRCm38) followed by nUMI and barcode counting, constructing the nUMI count matrices. The quality of the sequence alignment was checked using MultiQC (v1.9)^54^. nUMI matrices were filtered, centered, normalized, clustered and marker genes extracted using Scanpy (v1.8.1)^55^. Due to strong batch effects observed between the biological replicates, replicates were not merged but analyzed independently. Data for “control replicate 2” from E14.5 mice is shown. Leiden algorithm was used to cluster cells at different resolutions between 0.1 and 1 at the interval of 0.1. Resolution 0.8 is displayed. Marker genes were obtained with default parameter of the Scanpy’s find-marker function. Marker gene lists were automatically selected using rank gene groups (-rgg option) in scanpy functions. Gene Ontology enrichment was performed using hypergeometric test implemented in gseapy (v0.10.5)^56^.

### Atomic force microscopy (AFM)

AFM measurements of hair follicle basement membranes were performed on freshly cut 16 μm cryosections using JPK NanoWizard 2 (Bruker Nano) atomic force microscope mounted on an Olympus IX73 inverted fluorescent microscope (Olympus) and operated via JPK SPM Control Software v.5. Cryosections were equilibrated in PBS supplemented with protease inhibitors and measurements were performed within 20 minutes of thawing the samples. Triangular non-conductive Silicon Nitride cantilevers (MLCT, Bruker Daltonics) with a nominal spring constant of 0.07 Nm^-1^ were used for the nanoindentation experiments of the apical surface of cells and nuclei. For all indentation experiments, forces of up to 3nN were applied, and the velocities of cantilever approach and retraction were kept constant at 2μm^*s*-1^ ensuring an indentation depth of 500 nm. All analyses were performed with JPK Data Processing Software (Bruker Nano). Prior to fitting the Hertz model corrected by the tip geometry to obtain Young’s Modulus (Poisson’s ratio of 0.5), the offset was removed from the baseline, contact point was identified, and cantilever bending was subtracted from all force curves.

## ACKNOWLEDGEMENTS

We thank Carien M. Niessen and Leah C. Biggs for advice and critical reading of the manuscript, Anu M. Luoto, Hanne Ahola, and Claudia Ortmeier for expert technical assistance, Michele M. Nava for help with the AFM analyses, Fabien Bertillot for help with image quantification, the Max Planck Institute BioOptics and Biomedicum Helsinki Imaging Unit for imaging support and HiLIFE Laboratory Animal Centre Core Facility, University of Helsinki, for support with animal experiments. This work was supported by the Sigrid Juselius Foundation, Helsinki Institute of Life Science, Wihuri Research Institute, European Research Council (ERC) under the European Union’s Horizon 2020 research and innovation programme (grant agreement 770877 - STEMpop), and Academy of Finland Center of Excellence Barrier-Force (all to SAW). CV is the recipient of the Marie Curie fellowship H2020-MSCA-IF-101032331. YAM was the recipient of the EMBO Long-Term fellowship ALTF 728-2017 and Human Frontier Science Program fellowship LT000861/2018. MLMa and ELK acknowledge support from NIH R01HD099031 and Simons Foundation (454947 and 446222).

---

**Supplementary Figure 1.**
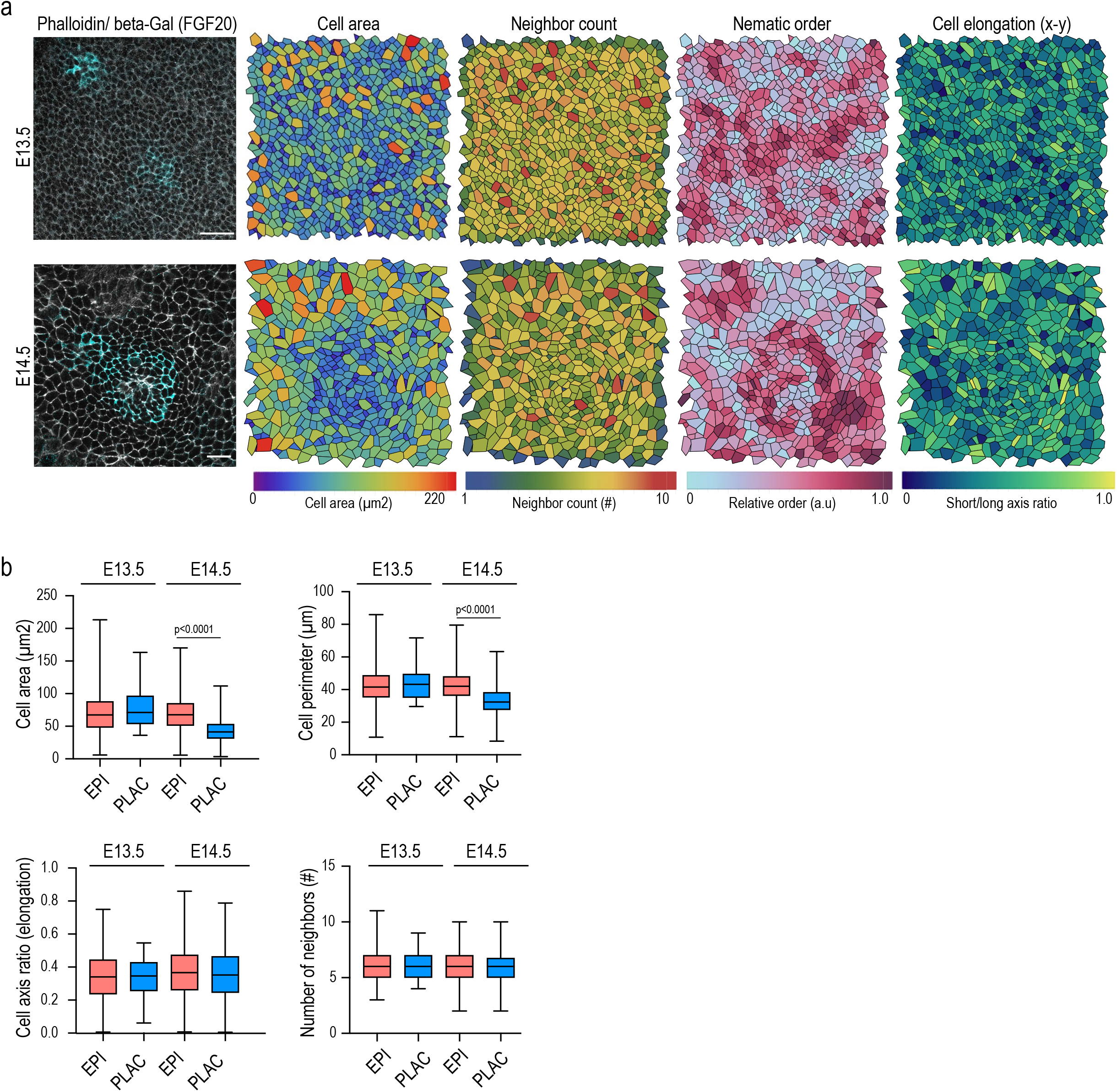
**(a)** Representative raw images of E13.5 and E14.5 cell membranes and FGF20-expressing (β-gal-stained) placode cells (left panels) and quantification of cell area, neighbor count, nematic order and cell elongation (in x-y) from segmented images. Scale bars 50 μm. Images representative of 3 mice/group. **(b)** Quantification of morphological parameters from images in (a). No substantial differences in cell morphology between placode and epidermis are observed at E13.5 (n>300 cells pooled across 3 mice /group).

**Supplementary Figure 2.**
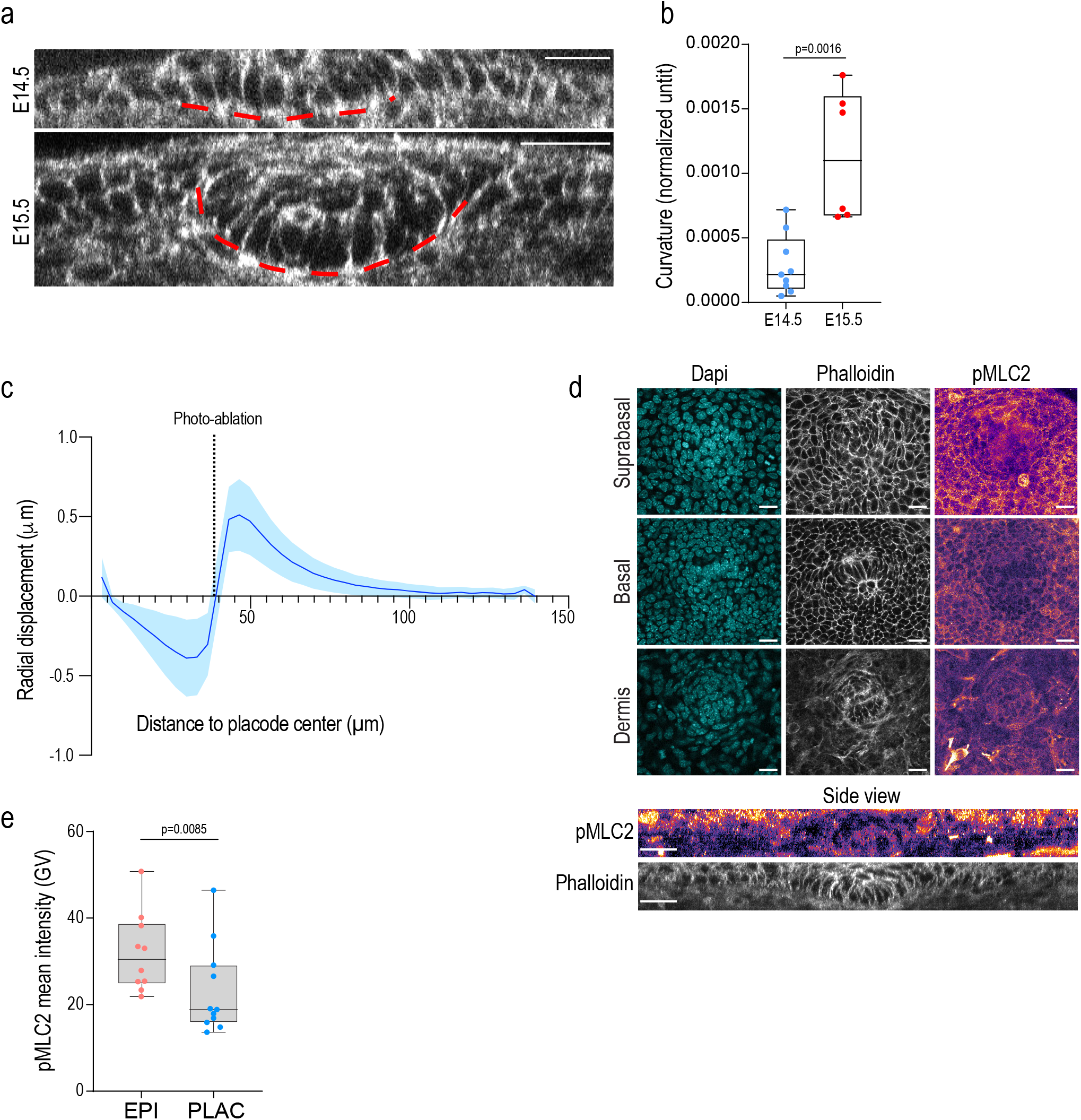
**(a, b)** Representative images (a) and quantification (b) of placode curvature at E14.5 and E15.5 (n= 9 (E14.5) or 6 (E15.5) placodes pooled across 3 embryos/ stage; Mann-Whitney). **(c)** Radial recoil as a function of distance to placode center from cuts between placode and epidermis (n=4 mice with 8-10 placodes/mouse). **(d)** Representative DAPI, phalloidin and pMLC2-stained images from E15.5 mouse skin whole-mount. Scale bars 20 μm. **(e)** Quantification of images from (d) shows a decrease in pMLC2 intensity in placode at E15 (n=10 placodes from 3 explants/condition; paired t-test).

**Supplementary Figure 3.**
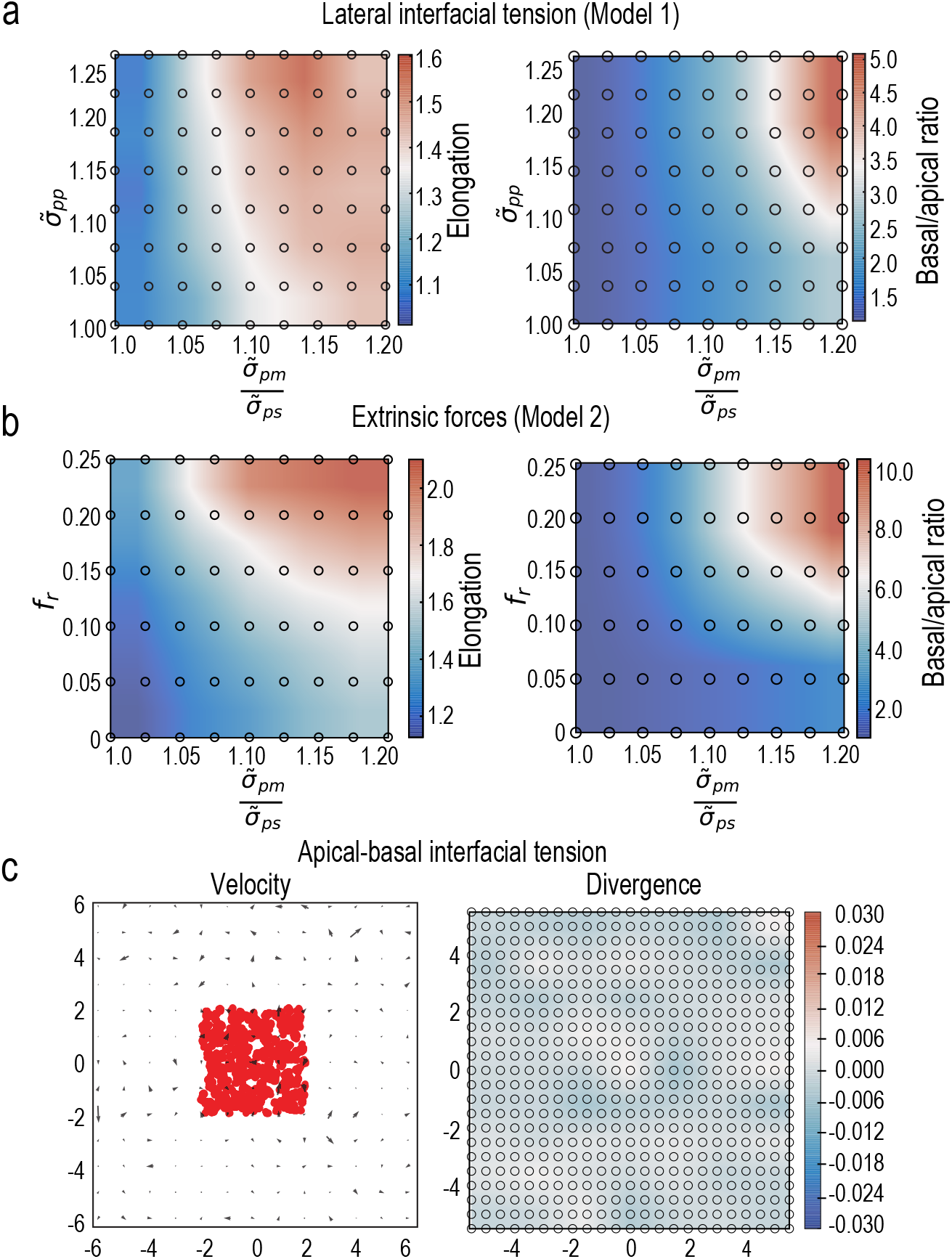
**(a, b)** Phase diagrams of cell elongation or changes in basal to apical surface length induced by interaction of lateral wetting coefficient (a) or extrinsic lateral forces (b) with varying basal to apical wetting coefficient. **(c)** Velocity map and mean divergence heatmap of model simulations applying high basal to apical wetting coefficient.

**Supplementary Figure 4.**
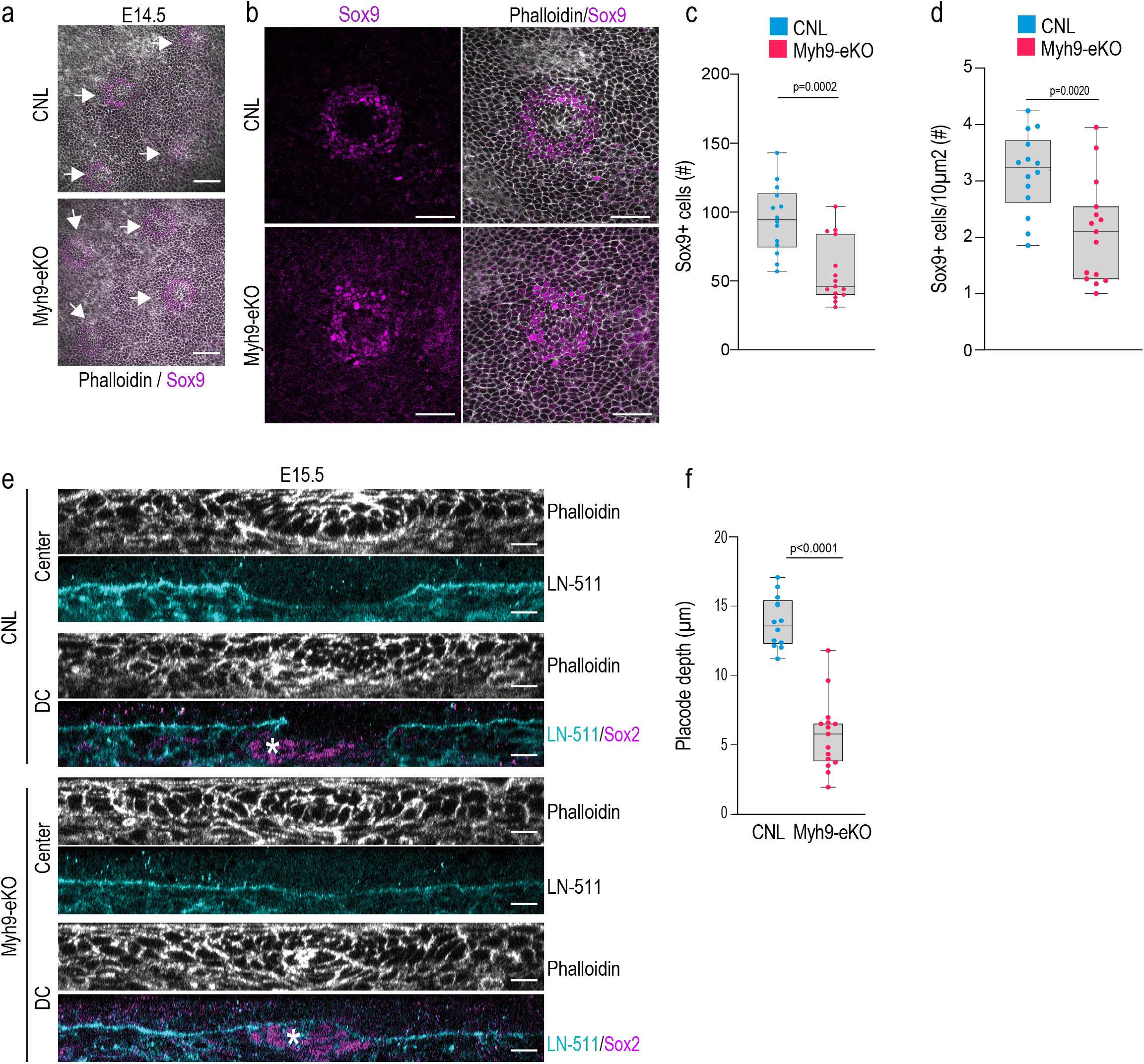
**(a)** Representative top views from basal layer of E14.5 CNL and Myh9-eKO mouse epidermis. White arrows mark Sox9-positive placodes. Scale bars 50 μm. Images representative of 4 mice/group. **(b-d)** Representative images (b) and quantification (c, d) of Sox9 pattern of expression from E15.5 CNL and Myh9-eKO mouse epidermis. Note reduced number of Sox9-positive cells (c) and local density (d) in Myh9-eKO placodes. Scale bars 50 μm. (n=14 (CNL) and 15 (Myh9-eKO) placodes from 3 mice/group. (c) Mann-Whitney; (d) Student’s t-test. **(e)** Phalloidin, LN-511 and Sox2-stained sagittal cross sections of CNL and Myh9-eKO mouse epidermis at E15.5. Sections from placode center and dermal condensate (DC; asterisk) are shown. Note position of DC within the placode in Myh9-eKO. Scale bars 10 μm. Images representative of 3 mice/group. **(f)** Quantifications of placode depth from CNL and Myh9-eKO mice at E15.5 shows reduced placode depth in Myh9-eKO mice (n=12 (CNL) and 15 (Myh9-eKO) placodes from 3 mice (left); Mann-Whitney).

**Supplementary Figure 5.**
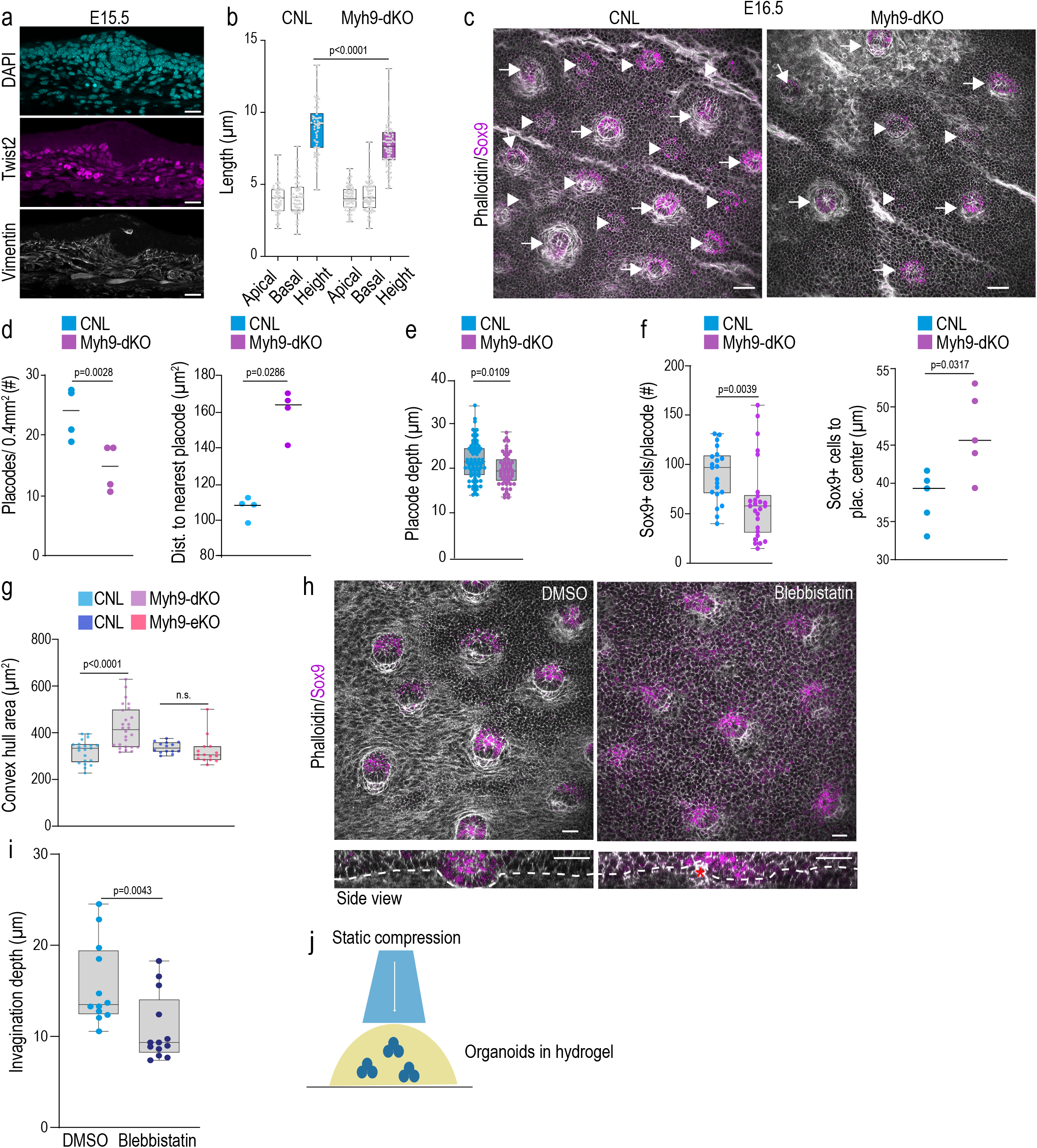
**(a)** Representative skin sections from E15.5 epidermis stained for DAPI, Vimentin and Twist-2 shows Twist-2-positive fibroblasts positioned below to the epidermis. Scale bars 20 μm. **(b)** Quantifications of placode cell surface lengths from CNL and Myh9-dKO mice at E14.5 shows reduced basal placode cell elongation in Myh9-dKO mice (n=113 (CNL) and 126 (Myh9dKO) placode cells pooled across 5 mice/group; one-way ANOVA). **(c-e)** Representative Sox9 and phalloidin-stained top views (c) and quantification (d, e, f) of skin wholemounts from CNL and Myh9-dKO mouse epidermis at E16.5 shows reduced number of second wave placodes (arrowheads) and invagination in myosin-deficient mice. Arrows mark first wave placodes. Scale bars 50 μm (n= 5 mice /group;Mann-Whitney, left panels; n>54 placodes pooled across 5 mice/group; Student’s t-test, right panel). **(f)** Quantification of the number of Sox9-postive cells (left) and the mean distance of Sox9-positive cells from the center of the placode (right) in skin whole mounts from CNL and MYh9-dKO epidermis at E14.5. (n=21 (CNL) and 26 (Myh9-dKO) placodes from 5 mice/group; Mann-Whitney; left and n=5 mice/group; Mann-Whitney; right). **(g)** Quantification of the minimum area covering Sox9-positive cells (convex hull area) shows a wider distribution of Sox9 expression in Myh9-dKO in contrast to CNL and Myh9-eKO (n=21 (CNL), 26 (Myh9-dKO), 14(CNL) and 15 (Myh9-eKO) placodes pooled across 3 mice (Myh9-eKO / CNL) or 5 mice (Myh9-dKO / CNL); one-way ANOVA). **(h, i)** Representative phalloidin-stained images (h) and quantification (i) of skin explants treated with Blebbistatin for 24h starting at E14.5. Note attenuated placode invagination and partial indentation of placode by the DC (red asterisk) in blebbistatin-treated explants. Scale bars 30 μm (n=12 (DMSO) and 13 (Blebbistatin) placodes from 3 mice; Mann-Whitney). **(j)** Schematic representation of embryonic epidermal organoid compression experiments.

**Supplementary Figure 6.**
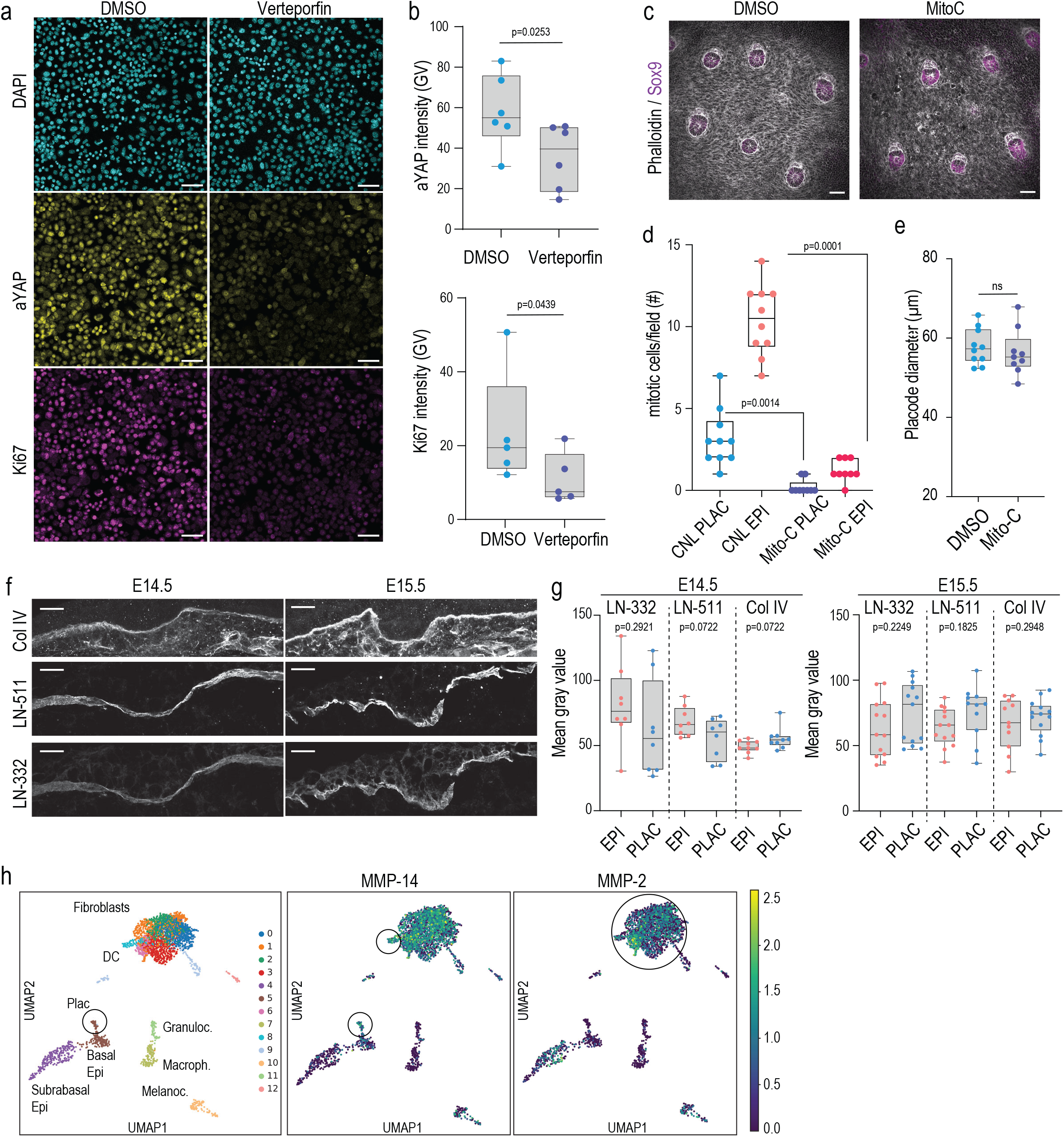
**(a, b)** Representative images (a) and quantification (b) of active YAP (aYAP; top) and proliferation (Ki67; bottom) in isolated epidermal stem cells treated with YAP inhibror Verteporfin (10 μM, 6 h). Scale bars 50 μm (n= 6 independent experiments, paired t-test). **(c)** Representative top views of Phalloidin/Sox9-stained Mito-C- and DMSO-treated control explants isolated from E14.5 mice. Scale bars 50 μm. Images representative of 3 mice/group. **(d)** Quantification of mitotic cells in placodes (PLAC) and interfollicular epidermis (EPI) of skin explants from E14.5 mice treated Mito-C. Note reduced mitoses in Mito-C-treated explants (n=10 (CNL), 9 (Mito-C) placodes (PLAC) or surrounding epidermis (EPI) from 3 mice/group; Kruskal-Wallis/Dunn’s. **(e)** Quantification of placode diameter in explants from E14.5 mice treated Mito-C (n=10 (CNL), 9 (Mito-C) placodes from 3 mice/group; Student’s t-test). **(f)** Collagen IV (Col IV), laminin (LN)-511, and LN-332-stained cross sections of E14.5 and E15.5 epidermis. Scale bars 10 μm. Images representative of 3 mice/group. **(g)** Quantification of images from (f) show no substantial differences in Col IV or LN-332/511 intensity between epidermis (EPI) and placode (PLAC) (pooled across 3 mice/group). **(h)** UMAP projections of single cell RNA seq data from E14.5 skin. Note MMP-14 expression in placode and fibroblast cells and MMP-2 expression in fibroblasts (black circles).

**Supplementary Table 1.**
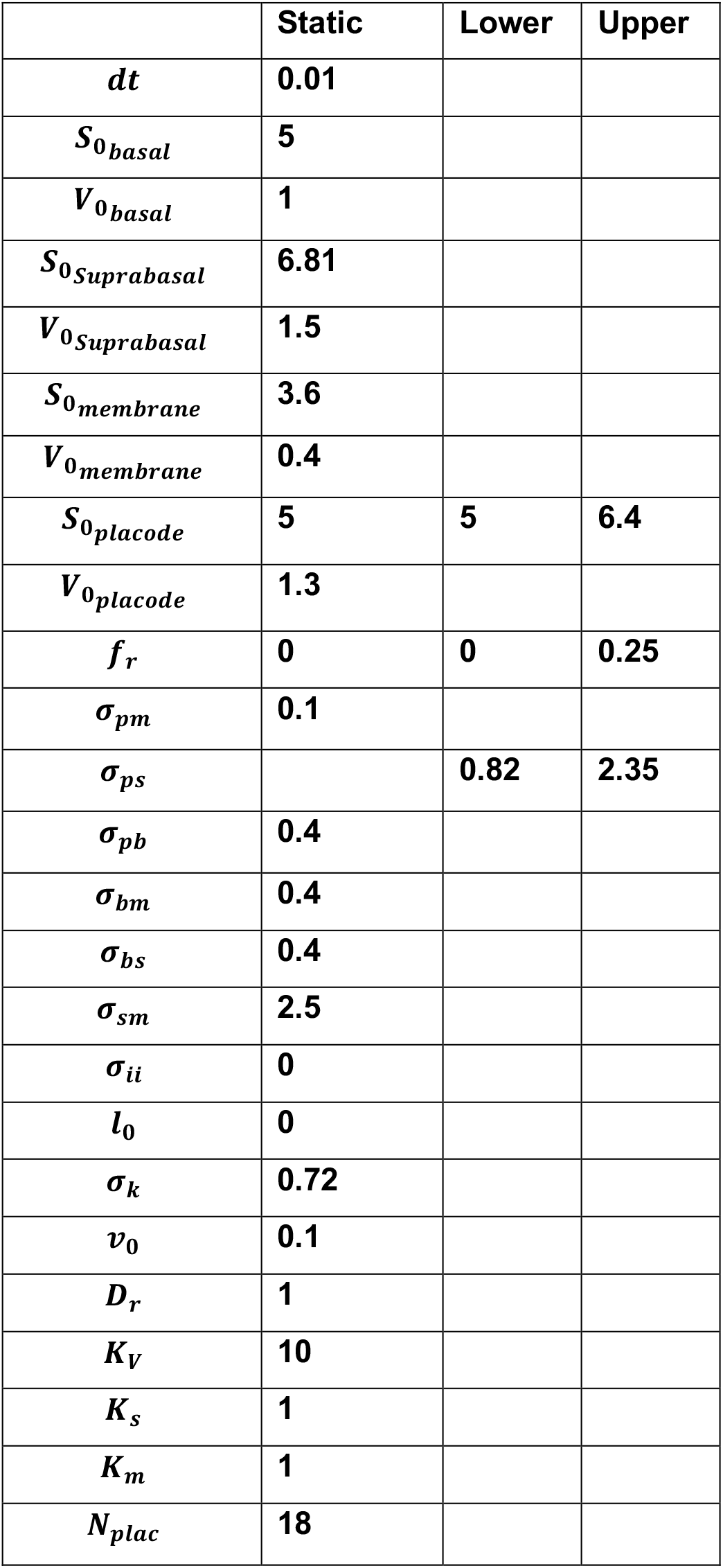
Parameter selection for 3D vertex model

|  | Static | Lower | Upper |
| --- | --- | --- | --- |
| $dt$ | 0.01 | | |
| $S_{0basal}$ | 5 | | |
| $V_{0basal}$ | 1 | | |
| $S_{0Suprabasal}$ | 6.81 | | |
| $V_{0Suprabasal}$ | 1.5 | | |
| $S_{0membrane}$ | 3.6 | | |
| $V_{0membrane}$ | 0.4 | | |
| $S_{0placode}$ | 5 | 5 | 6.4 |
| $V_{0placode}$ | 1.3 | | |
| $f_r$ | 0 | 0 | 0.25 |
| $\sigma_{pm}$ | 0.1 | | |
| $\sigma_{ps}$ | | 0.82 | 2.35 |
| $\sigma_{pb}$ | 0.4 | | |
| $\sigma_{bm}$ | 0.4 | | |
| $\sigma_{bs}$ | 0.4 | | |
| $\sigma_{sm}$ | 2.5 | | |
| $\sigma_{ii}$ | 0 | | |
| $l_0$ | 0 | | |
| $\sigma_k$ | 0.72 | | |
| $v_0$ | 0.1 | | |
| $D_r$ | 1 | | |
| $K_V$ | 10 | | |
| $K_s$ | 1 | | |
| $K_m$ | 1 | | |
| $N_{plac}$ | 18 | | |

